# AAA+ protease-adaptor structures reveal altered conformations and ring specialization

**DOI:** 10.1101/2022.02.04.479121

**Authors:** Sora Kim, Xue Fei, Robert T. Sauer, Tania A. Baker

## Abstract

ClpAP, a two-ring AAA+ protease, degrades N-end-rule proteins bound by the ClpS adaptor. Here, we present high-resolution cryo-EM structures of ClpAPS complexes showing how ClpA pore loops interact with the ClpS N-terminal extension (NTE), which is normally intrinsically disordered. In two structural classes, the NTE is bound by a spiral of pore-1 and pore-2 loops in a manner similar to substrate-polypeptide binding by many AAA+ unfoldases. Kinetic studies reveal that pore-2 loops of the ClpA D1 ring catalyze protein remodeling required for substrate delivery by ClpS. In a third class, D2 pore-1 loops are rotated and tucked away from the channel, and do not bind the NTE, demonstrating asymmetry in engagement by the D1 and D2 rings. These studies demonstrate new structures and functions for key AAA+ elements. In addition to ClpAPS delivery, pore-loop tucking may be used broadly by AAA+ unfoldases, for example during enzyme pausing/unloading.

## INTRODUCTION

Regulated proteolysis by energy-dependent AAA+ (ATPases associated with diverse cellular <u>a</u>ctivities) proteases maintains cellular homeostasis of proteins in all organisms. AAA+ proteases harness the energy of ATP hydrolysis to degrade regulatory proteins and proteins that are damaged, misfolded, or no longer needed (Sauer and Baker, 2011). AAA+ proteases, such as ClpAP, consist of a hexameric AAA+ unfoldase (*e.g*. ClpA_6_) and a self-compartmentalized peptidase (*e.g*. ClpP_14_). In the recognition step of degradation, a peptide sequence (degron) in a protein substrate is engaged by pore loops lining the axial channel of the AAA+ hexamer. Through conformational changes powered by ATP-hydrolysis cycles, native structure in the bound substrate is unfolded and processively translocated through the channel and into the peptidase chamber, where the polypeptide is cleaved into fragments. In addition to binding and engaging degrons directly, AAA+ proteases interact with adaptor proteins that modify their substrate specificity, often by tightening the affinity of the enzyme•substrate complex (Sauer and Baker, 2011; Olivares et al., 2015; Mahmoud and Chien, 2018).

Prokaryotes and eukaryotes use the N-end-rule pathway to target proteins bearing specific N-terminal residues (called N-degrons) for rapid degradation (Varshavsky, 2019). In *Escherichia coli*, the ClpS adaptor promotes ClpAP degradation of proteins containing Leu, Phe, Tyr, or Trp residues at the N-terminus (Tobias et al., 1991; Dougan et al., 2002; Zeth et al., 2002a, 2002b; Erbse et al., 2006; Varshavsky, 2019). The ClpS protein (~10 kDa) docks with the N-terminal domain of ClpA and contains a hydrophobic pocket that binds the N-end-rule residue (Zeth et al., 2002a, 2002b; Wang et al., 2008a; Román-Hernández et al., 2009). ClpS functions as a specificity switch for ClpAP, promoting degradation of N-degron substrates while inhibiting degradation of ssrA-tagged and related substrates (Dougan et al., 2002; Guo et al., 2002; Erbse et al., 2006; Wang et al., 2008a). Unlike some adaptors that simply act as molecular matchmakers between the substrate and enzyme, ClpS also modulates the rate of ATP hydrolysis and substrate unfolding by ClpA. Interestingly, ClpS is proposed to interact with ClpA as a ‘pseudo-substrate’ (Dougan et al., 2002; De Donatis et al., 2010; Román-Hernández et al., 2011; Rivera-Rivera et al., 2014; Torres-Delgado et al., 2020). Specifically, the N-terminal extension (NTE) of free ClpS is exposed as an unstructured peptide, mimicking a degron. The NTE is poorly conserved among orthologs, with the exception of a short junction sequence adjacent to the ClpS core that typically contains a few tandem prolines (Hou et al., 2008; Román-Hernández et al., 2011). During ClpS-assisted degradation, a ClpS•N-degron substrate complex initially binds to ClpA. Subsequently, the N-degron substrate is transferred to ClpA for degradation and ClpS escapes destruction by mechanisms that are poorly understood.

Each ClpA subunit has two AAA+ modules, called D1 and D2, that associate in the hexamer to form two stacked rings (Grimaud et al., 1998). The D1 and D2 modules belong to different AAA+ subfamilies and have distinct biochemical functions (Erzberger and Berger, 2006). The D2 ring, a member of the HCLR AAA+ clade, is the principal ATPase motor responsible for unfolding and translocating substrates, including proteins with high thermodynamic stabilities (Kress et al., 2009; Kotamarthi et al., 2020; Zuromski et al., 2020). In contrast, the D1 ring, a classic AAA+ clade member, assists the D2 ring as an auxiliary motor, improves enzyme processivity, and plays a major role in substrate recognition (Kress et al., 2009; Kotamarthi et al., 2020; Zuromski et al., 2020, 2021). ClpS differentially regulates the activities of the D1 and D2 rings (Kress et al., 2009; Zuromski et al., 2021) via interactions of its NTE that we characterize here. Previous cryo-EM structures of ClpAP elucidate how the axial channel of the D1 and D2 rings engages the polypeptide of a directly recognized substrate (Lopez et al., 2020). Pore-1 and pore-2 loops in both rings form spiral-staircase-like arrangements that bind the substrate polypeptide, in a similar manner to those in structures of other double-ring AAA+ enzymes, such as Hsp104, ClpB, Cdc48/p97, and NSF (Zhao et al., 2016; Deville et al., 2017; Gates et al., 2017; White et al., 2018; Yu et al., 2018; Cooney et al., 2019; Lo et al., 2019; Rizo et al., 2019; Twomey et al., 2019). However, these previous structures do not provide insight into the distinct, specialized functions of each AAA+ ring of ClpA or the mechanism of ClpS-assisted degradation of N-degron substrates.

To characterize ring specialization and ClpS-ClpA collaboration, we solved cryo-EM structures of ClpAPS complexes that show how the normally disordered ClpS NTE assumes an extended conformation when bound in the ClpA axial channel. These structures reveal marked conformational differences from prior ClpAP structures (Lopez et al., 2020). We identify multiple conformations of ClpS-bound ClpA, including an arrangement in which the pore-1 loops of the D2 ring are tucked-in and face away from the channel, allowing only the D1 ring to interact strongly with the ClpS NTE. Mutagenesis and biochemical experiments establish that the pore-2 loops of the ClpA D1 ring are essential for ClpS delivery of an N-degron substrate but contribute little to docking of ClpS with ClpA. Our results demonstrate structural and functional plasticity among the ClpA pore loops, provide a structural basis for the functions of ClpS during N-degron substrate degradation, and contribute more broadly to understanding the operational modes available to AAA+ enzymes as they perform diverse biological processes.

## RESULTS

### Distinct conformations of ClpAPS delivery complexes

We used size-exclusion chromatography in the presence of ATPγS to purify a complex of ClpA, ClpP, ClpS, and the N-end-rule substrate YLFVQELA-GFP (**Figure 1A-B**). Based on SDS-PAGE, the YLFVQELA-GFP substrate appeared to be sub-stoichiometric compared to ClpS (**Figure 1B**). Because ATPγS does not support degradation (Thompson et al., 1994; Hoskins et al., 1998; Ishikawa et al., 2001; Effantin et al., 2010; Miller and Lucius, 2014; Lopez et al., 2020), these complexes should represent early stages in ClpS-mediated delivery of N-degron substrates.

**Figure 1.**
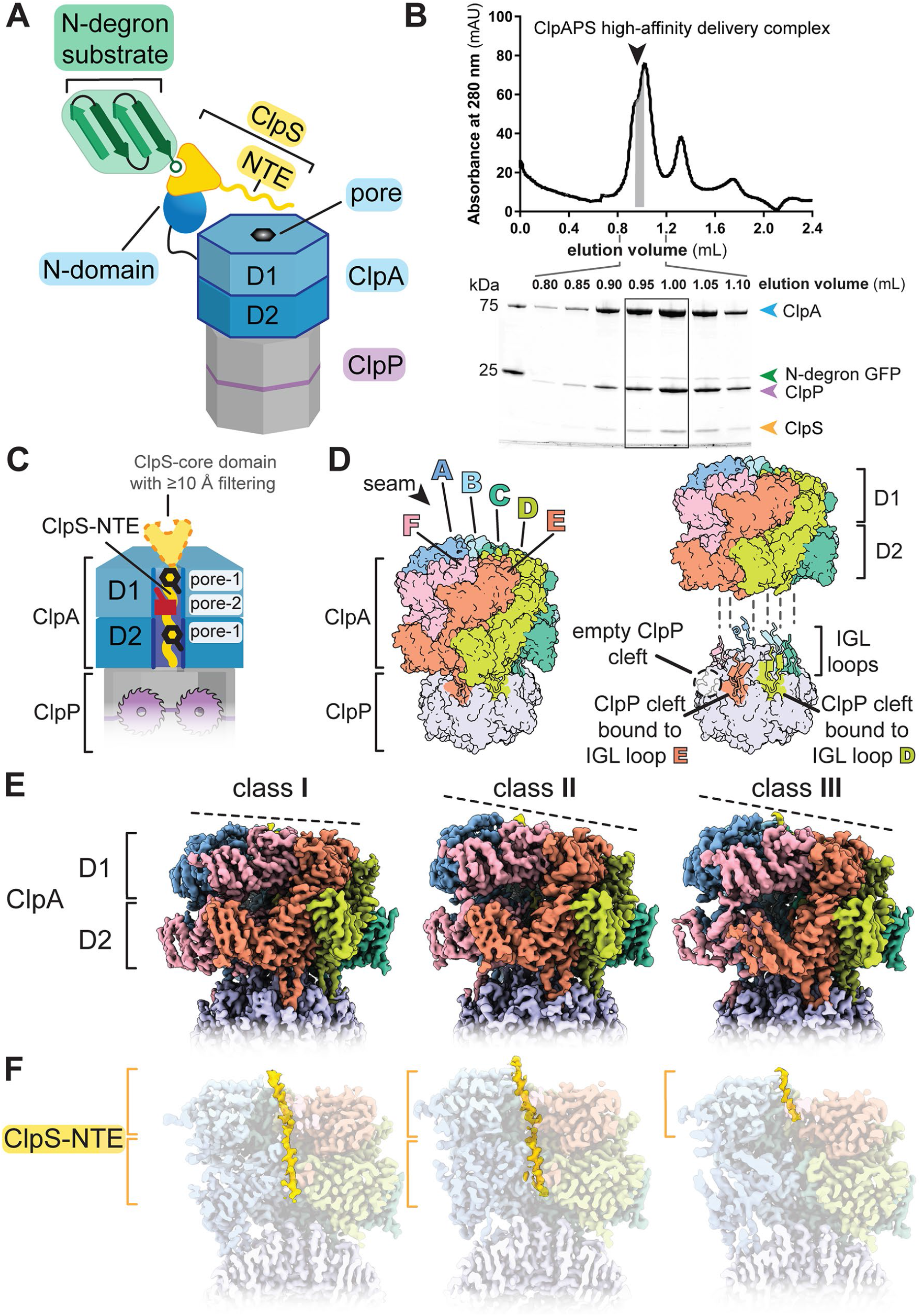
Architectures of ClpS-bound ClpAP. (A) Cartoon of ClpS delivery of an N-degron substrate for ClpAP degradation. Native ClpS (yellow wedge) binds to the N-degron substrate (green) and also binds to an N-terminal domain of ClpA (blue oval) (B) Size-exclusion chromatography (top panel) of a complex of ClpA, ClpP, ClpS, and YLFVQELA-GFP (an N-degron substrate) assayed by SDS-PAGE (bottom panel). Gray shaded area and boxed area indicate the fractions pooled for cryo-EM. (C) Cartoon of pore loops that interact with the NTE and proteins resolved in cryo-EM structures. (D) (left panel) ClpA subunit nomenclature in right-hand spiral hexamer, where the seam interface is between the lowest (F) subunit and the highest (A) subunit. The ClpA hexamer docks into clefts in the ClpP_7_ ring via IGL loops (right panel). The empty ClpP cleft is located between the clefts occupied by subunits E and F. (E) Side views of the cryo-EM maps of classes **I**, **II**_**c**_, and **III**_**b**_. The dashed line indicates the relative height of ClpA subunits within the spiral. (F) Cutaway views of panel E showing density for the ClpS NTE colored yellow.

Following single-particle cryo-EM analyses (**Figure S1**), three-dimensional (3D) classification and reconstruction using RELION-3 yielded six density maps (3.22–3.38 Å), representing three general structural classes (**I**, **II**, and **III**) with the latter classes being subdivided into **II**_**a**_/**II**_**b**_/**II**_**c**_ or **III**_**a**_/**III**_**b**_ subclasses (**Table 1**; **Figure S2**). In low-pass filtered maps, the ClpS core domain (res. 27–106) could be docked into each of the six maps (**Figure S3**). In unfiltered maps, there was good density for all or part of the NTE of ClpS, for the D1 and D2 rings of ClpA, and for ClpP (**Figure 1C**). There was no substantial density for the core domain of ClpS, the N-domains of ClpA, or YLFVQELA-GFP, suggesting that these domains/proteins are not present in fixed conformations relative to the remaining parts of the complex or are potentially absent (YLFVQELA-GFP).

**Table 1.** Cryo-EM data collection, processing, model building, and validation statistics.

| **class name** | **A** | **B** |  |  | **C** |  |
| --- | --- | --- | --- | --- | --- | --- |
| PDB ID | XXX | XXX | XXX | XXX | XXX | XXX |
| EMDB ID | XXX | XXX | XXX | XXX | XXX | XXX |
| **data collection and processing** |  |  |  |  |  |  |
| microscope | Talos Arctica |  |  |  |  |  |
| camera | K3 |  |  |  |  |  |
| magnification | 45,000X |  |  |  |  |  |
| voltage (kV) | 200 |  |  |  |  |  |
| total electron dose (e-/Å<sup>2</sup>) | 34.71 |  |  |  |  |  |
| defocus range (μm) | -0.5 to -2.5 |  |  |  |  |  |
| pixel size (Å) | 0.435 |  |  |  |  |  |
| micrographs collected | 9169 |  |  |  |  |  |
| initial particles | 1,043,033 |  |  |  |  |  |
| final particles | 51,750 | 156,677 | 43,431 | 37,530 | 37,885 | 31,453 |
| symmetry | C1 | C1 | C1 | C1 | C1 | C1 |
| map resolution (Å) at 0.143 FSC threshold | 3.24 | 3.38 | 3.33 | 3.24 | 3.22 | 3.26 |
| **model composition** |  |  |  |  |  |  |
| non-hydrogen atoms | 38021 | 37900 | 37776 | 38131 | 37880 | 38006 |
| protein residues | 4820 | 4805 | 4792 | 4837 | 4819 | 4823 |
| nucleotides | 12 | 11 | 10 | 12 | 12 | 12 |
| **refinement** |  |  |  |  |  |  |
| map-model CC | 0.80 | 0.77 | 0.75 | 0.81 | 0.81 | 0.82 |
| RMSD bond lengths (Å) | 0.005 | 0.005 | 0.005 | 0.005 | 0.006 | 0.006 |
| RMSD bond angles (°) | 1.115 | 1.117 | 1.100 | 1.095 | 1.156 | 1.142 |
| **validation** |  |  |  |  |  |  |
| MolProbity score | 1.09 | 1.06 | 1.06 | 1.07 | 1.14 | 1.10 |
| clash score | 2.98 | 2.74 | 2.74 | 2.78 | 3.48 | 3.05 |
| Cβ deviation (%) | 0.00 | 0.00 | 0.00 | 0.00 | 0.00 | 0.00 |
| rotamer outliers (%) | 0.00 | 0.00 | 0.02 | 0.00 | 0.00 | 0.00 |
| Ramachandran favored (%) | 99.94 | 99.73 | 99.75 | 99.88 | 99.77 | 99.50 |
| Ramachandran disallowed (%) | 0.00 | 0.00 | 0.00 | 0.00 | 0.00 | 0.00 |

In our structures, the six subunits of the ClpA hexamer, which we label A through F (clockwise direction with subunit F at the bottom), formed a shallow spiral, as expected from prior cryo-EM structures (Lopez et al., 2020). Six flexible IGL loops (res. 610–628) in each ClpA hexamer docked into clefts in the heptameric ring of ClpP, leaving one empty cleft (**Figure 1D**). Differences between classes **I**, **II**, and **III** include the relative positions of subunits in the ClpA spiral, density for the ClpS NTE in the ClpA channel, and changes within individual ClpA subunits (**Figure 1E-F**; **Figure 2**). For example, density for the ClpS NTE was present in both the D1 and D2 rings of ClpA in classes **I** and **II**, but only in the D1 ring of class **III**(**Figure 1F**). In classes **II** and **III**, the relative height of ClpA subunits in the spiral was A (highest) > B > C > D > E > F (lowest), whereas in class-**I** subunit B was higher than subunit A, resulting in a more planar D1 ring (**Figure 2A-B**). Additionally, in the D2 ring of class-**III** structures, the small AAA+ domain of subunit E swings outward, breaking the rigid-body interface with its large AAA+ domain neighbor from subunit F (**Figure 2C**). By contrast, structures of ClpAP with RepA-GFP and ATPγS (Lopez et al., 2020) did not display this feature, suggesting that it arises as a consequence of ClpS binding. The subclasses (**II**_**a**_/**II**_**b**_/**II**_**c**_ or **III**_**a**_/**III**_**b**_) differed from each other largely in the detailed interactions between ClpA and the ClpS NTE, the visibility of individual NTE residues, and the nucleotide occupancy of each ATPase site (ATPγS, ADP, or empty) (**Figure S4–S5**).

**Figure 2.**
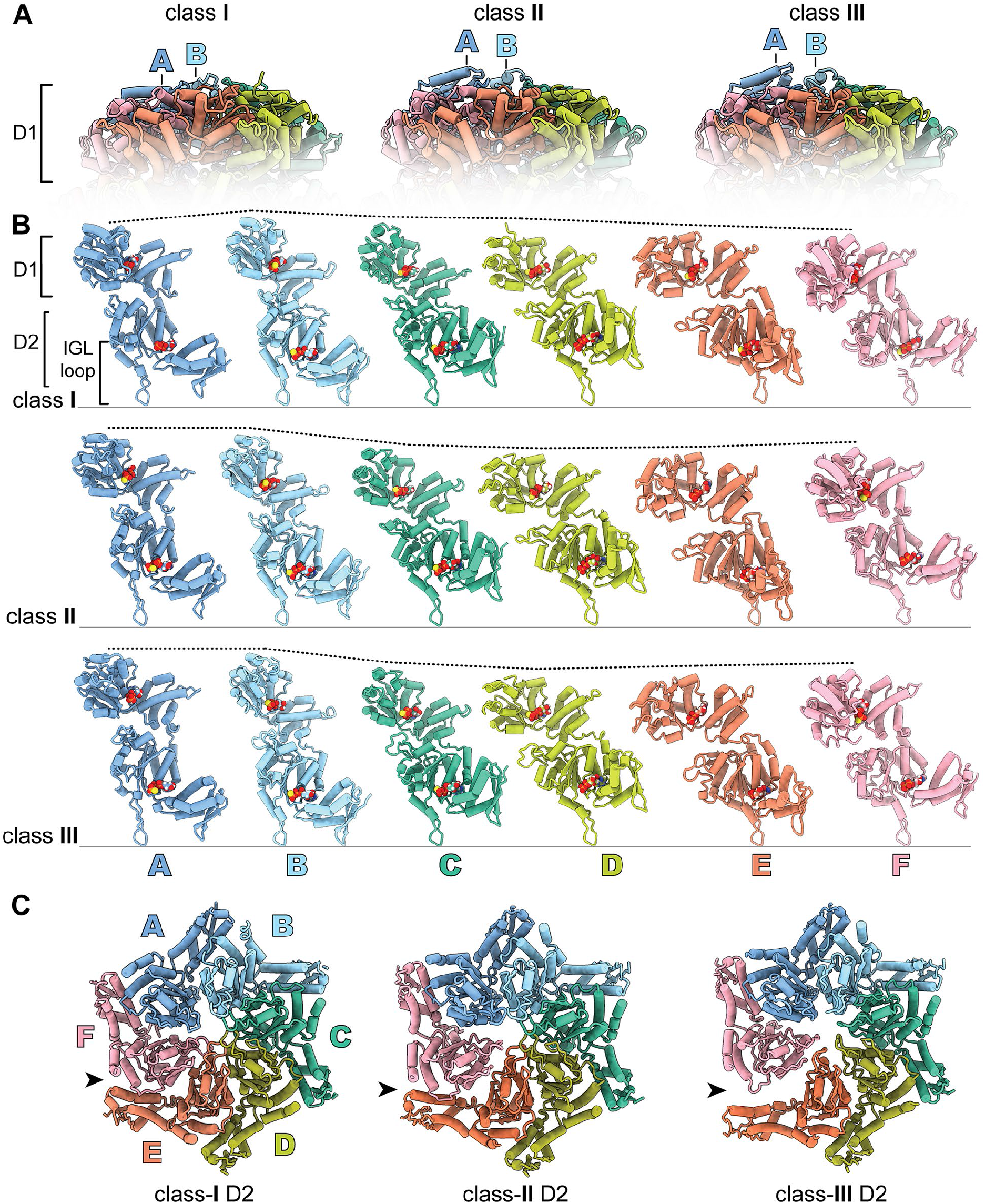
Conformational differences in ClpA subunits and hexamers. (A) Atomic models of class **I**, **II**_**c**_, and **III**_**a**_ D1 rings, showing position of subunits A and B. (B) Individual subunits of class **I**, **II**_**c**_, and **III**_**a**_ atomic models. The dashed line indicates the relative height of each subunit, following alignment to the bottom of the IGL loop. (C) D2 ring rigid-body interface between subunits E and F of classes **I**, **II**_**c**_, and **III**_**a**_. In class **III**_**a**_, the small AAA+ domain of subunit E in the D2 ring swings out and loses contact (arrow) with the neighboring large D2 AAA+ domain of subunit F.

### Binding of the ClpS NTE within the axial channel of ClpA

Each of our structures contained clear main-chain and side-chain density corresponding to all or part of the ClpS NTE (res. 2–26) in the ClpA channel (**Figure 3A**). The register of this ClpS peptide is very similar in each structure, with the C-terminal portion of the NTE (Pro^24^–Pro^25^–Ser^26^) near the top of the ClpA channel, and the N-terminal portion near the bottom of the channel in classes **I** and **II**. The ClpA channel is lined by the D1 K<u>Y</u>R pore-1 loops (res. 258–260) and pore-2 loops (res. 292–302) and by the D2 G<u>Y</u>VG pore-1 loops (res. 539–542) and pore-2 loops (res. 526–531). Pore-1 loops of AAA+ unfoldases and protein-remodeling machines contain a key, conserved aromatic side chain (usually tyrosine; underlined in K<u>Y</u>R and G<u>Y</u>VG) that contacts the substrate polypeptide in the channel and functions in the binding, unfolding, and translocation of target proteins (Schlieker et al., 2004; Weibezahn et al., 2004; Hinnerwisch et al., 2005a; Martin et al., 2008; Doyle et al., 2012; Iosefson et al., 2015; Lopez et al., 2020; Zuromski et al., 2021). The ClpS NTE was bound by many K<u>Y</u>R and G<u>Y</u>VG pore-1 loops and also by the D1 pore-2 loops of ClpA. Neighboring pore-1 loops interacted with two-residue segments of the NTE, as observed for substrate polypeptides bound to multiple AAA+ unfoldases and protein-remodeling machines (Puchades et al., 2020).

**Figure 3.**
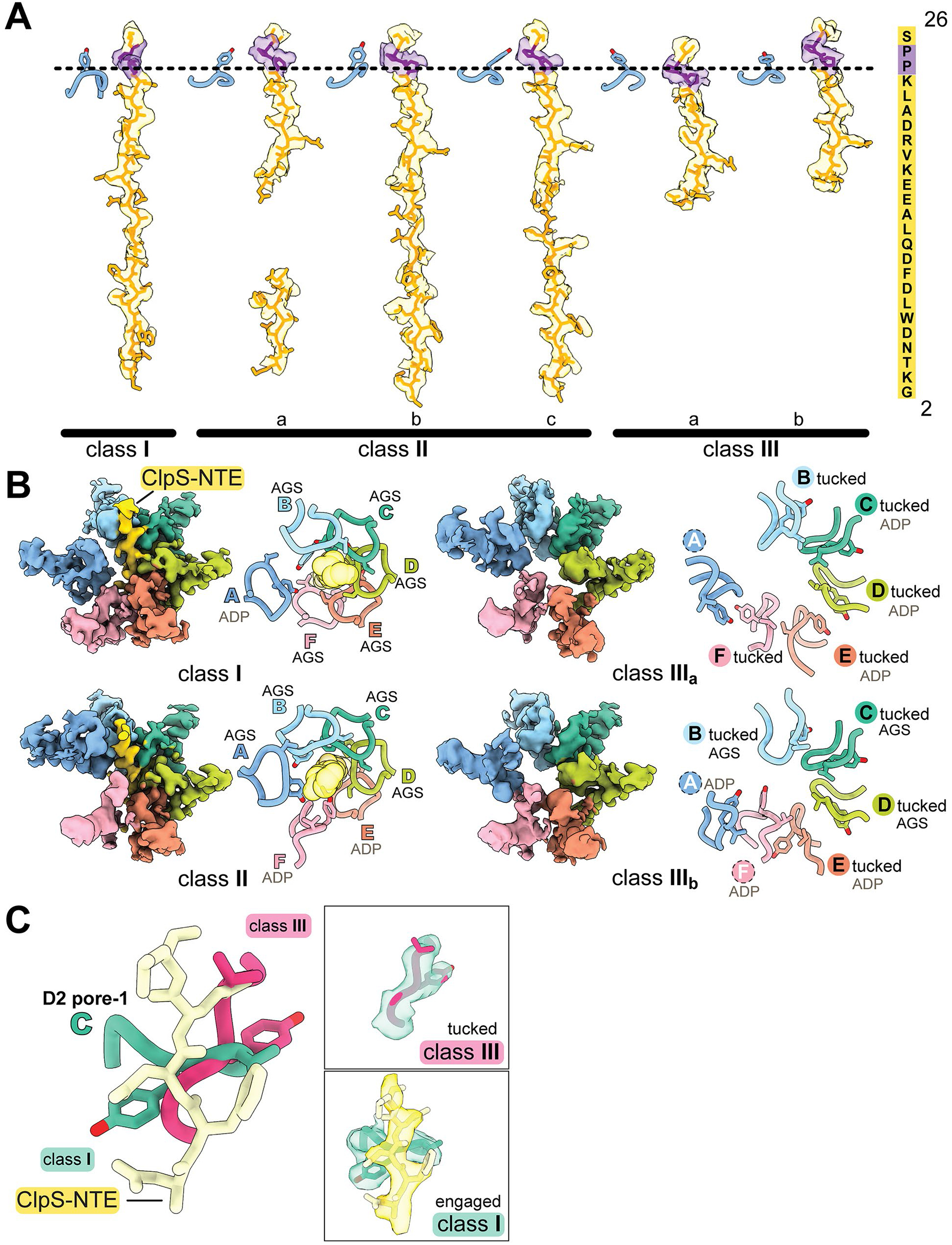
Conformations of the ClpS NTE and D2 pore-1 loops of ClpA. (A) Density of the ClpS NTE (transparent surface with modeled residues in sticks) in the ClpA axial channel in classes **I**, **II**, and **III**. Pro^24^ and Pro^25^ (colored purple) are part of the junction sequence between the NTE and the ClpS core domain. The ClpS NTE sequence is shown on the right. The D1 pore-1 loop of subunit A is shown as a reference point for the top of ClpA. (B) D2 ring ClpA pore-1 loops and the ClpS NTE in classes **I**, **II**_**c**_, and **III**_**a,b**_. The left panel in each structure depicts cryo-EM density for ClpA res. 528–555 and the ClpS NTE res. 2–15. The right panels are a zoomed-in view of the pore-1 loops (res. 538–542) and the NTE (transparent spheres), which is absent in class-**III**, in the atomic models. Subunit labels indicate nucleotide and interaction with the ClpS NTE. Labels in colored text denote NTE engagement; the dotted circle denotes lack of NTE engagement, with Tyr^540^ pointing towards the channel; labels in black text indicate the tucked conformation (Tyr^540^ away from the channel). (C) Atomic models (sticks) and density (transparent surfaces in boxed area) of the subunit C D2 pore-1 loop in classes **I**(green) or **III**_**b**_ (pink) and the class-**I** ClpS NTE (yellow). In class **III**, density for the NTE is not observed.

Despite this overall resemblance to substrate engagement, there were deviations in individual pore-1 loop interactions from those in prior structures of ClpAP and some Hsp100 family members. For example, the D2 G<u>Y</u>VG pore-1 loops of all six ClpA subunits contacted the ClpS NTE in classes **I** and **II**_**c**_(**Figure S5**), whereas previous ClpAP structures and subclasses **II**_**a**_ and **II**_**b**_ show four or five engaged G<u>Y</u>VG loops (**Figure S6B**) (Lopez et al., 2020). The configuration of pore-1 loops in classes **I** and **II**_**c**_ was also different from an extended Hsp104•casein structure in which loops from both the top and bottom AAA+ rings of all six protomers contact substrate in a split ‘lock-washer’ conformation (Gates et al., 2017). In classes **I** and **II**_**c**_, we observed five bound and one unbound pore-1 loops in D1 and six bound pore-1 loops in D2, an arrangement found in the high-affinity state of *Mycobacterium tuberculosis* ClpB (Yu et al., 2018). In many AAA+ structures, only the pore loops of ATP-bound subunits contact substrate (Puchades et al., 2020). By contrast and as reported for ClpAP•substrate complexes (Lopez et al., 2020), the pattern of engaged vs. disengaged pore loops in our structures did not strictly correlate with the nucleotide present in the corresponding ATPase active site (**Figures S4–S5**). For instance, ADP is bound to the class-**II**_**c**_ D2 nucleotide sites in subunits E and F, but the G<u>Y</u>VG loops from these domains contact the NTE. The presence of 11 engaged pore-1 loops (five D1 and six D2) likely contributes to the high-affinity of ClpAPS•N-degron complexes assembled in ATPγS (Román-Hernández et al., 2011).

### Pore-1 loops of D2 ring rotate out of the channel to alter polypeptide contacts

In classes **I** and **II**, residues 2–15 of the ClpS NTE were built into density in the D2 portion of the channel, but this NTE region was not visible in class **III**, presumably as a consequence of its conformational heterogeneity. We infer that these NTE residues are within D2, as the more C-terminal NTE segment (res. 16–26) is bound by the D1 ring of class **III** in the same manner as in classes **I** and **II**. Thus, the two AAA+ rings of ClpA can differ in their engagement with the NTE, a feature not observed in substrate-bound ClpAP structures (Lopez et al., 2020).

This absence of density for the N-terminal portion of the NTE in class **III** correlated with distinct structural features within the axial channel. Most surprisingly, the D2 pore-1 loops in class **III** were rotated ~90° compared to their orientation in classes **I** and **II**, and the key Tyr^540^ side chains were tucked-in and turned away from the axial channel (**Figure 3B–C**; **Movie S1**). In both class-**III** subclasses, at least four of the six pore 1 loops were convincingly in this new tucked conformation.

In many AAA+ unfoldases and protein-remodeling machines, one or two pore-1 loops, usually at the top and bottom of the spiral, are disengaged from the substrate polypeptide as a result of translational displacement of the corresponding subunit(s) (Deville et al., 2017; Gates et al., 2017; Puchades et al., 2017, 2019, 2020; Ripstein et al., 2017, 2020; Yu et al., 2018; Dong et al., 2019; Han et al., 2019, 2020; Lo et al., 2019; Fei et al., 2020a, 2020b; Lopez et al., 2020). This ‘canonical’ disengaged state of pore-1 loops in one or two subunits is very different than the tucked and rotated orientations of the class-**III** D2 pore-1 loops, in which no interactions with the polypeptide in the channel were present in the D2 ring. Pore-1 tyrosine contacts with the polypeptide within the AAA+ channel are considered essential for substrate binding and translocation. Thus, rotation of most (or all) Tyr^540^ side chains in the class-**III** D2 ring is sufficient to explain the lack of initial engagement of the N-terminal segment of the NTE and/or loss of binding that may occur during ClpS-assisted degradation of N-degron substrates (see *Discussion*).

Three additional features of the class-**III** D2 ring are noteworthy. Coincident with the pore-1-loop rotation, the ClpA channel in the D2 ring of class **III** was wider than in classes **I** and **II** (**Figure 2C**). Second, as noted above, the D2 rigid-body interface between the small AAA+ domain of subunit E and its neighboring large AAA+ domain in subunit F was broken in class **III**. This rearrangement may facilitate the accompanying conformational changes that result in loss of NTE contacts by the D2 pore-1 loops. Finally, the D2 ring contained ADP in three adjacent subunits in class **III**, whereas classes **I** and **II** contained no more than two ADPs in the D2 ring (**Figure S4**). Thus, ClpA has the ability to bind all of a polypeptide in the axial channel tightly using pore loops in both rings or by disrupting coordinated activity of the D1 and D2 rings, to specifically bind only the C-terminal portion of this sequence within the D1 ring.

### Pore-2 loops in the D1 ring form a second network of NTE-engaging contacts

In addition to the pore-1 loop interactions described above, our structures show that at least four pore-2 loops (res. 292–302) in the D1 ring of ClpA contacted the ClpS NTE (**Figure 4**). In each subunit, these pore-2 contacts were positioned below the corresponding D1 K<u>Y</u>R contacts and were offset by ~60°. The Ala^295^-Ala^296^-Ser^297^ tripeptide (AAS) at the tip of the D1 pore-2 loops contacted the opposing face of the ClpS NTE compared to the contacts made by the D1 pore-1 loops (compare orientation of D1 pore-2 loops on left vs. D1 pore-1 loops on right side of channel in **Figure 4A**). In contrast to the well-defined K<u>Y</u>R motif in the D1 pore-1 loop, which is conserved among Hsp104/ClpABC protein-remodeling enzymes and contains the invariant aromatic residue present in all AAA+ unfoldases, the key residues and functions of the pore-2 loops have been poorly delineated to date (Puchades et al., 2020). Among ClpABC family members, the pore-2 loops are more variable in sequence and length (**Figure 4B**; **Figure S6A**).

**Figure 4.**
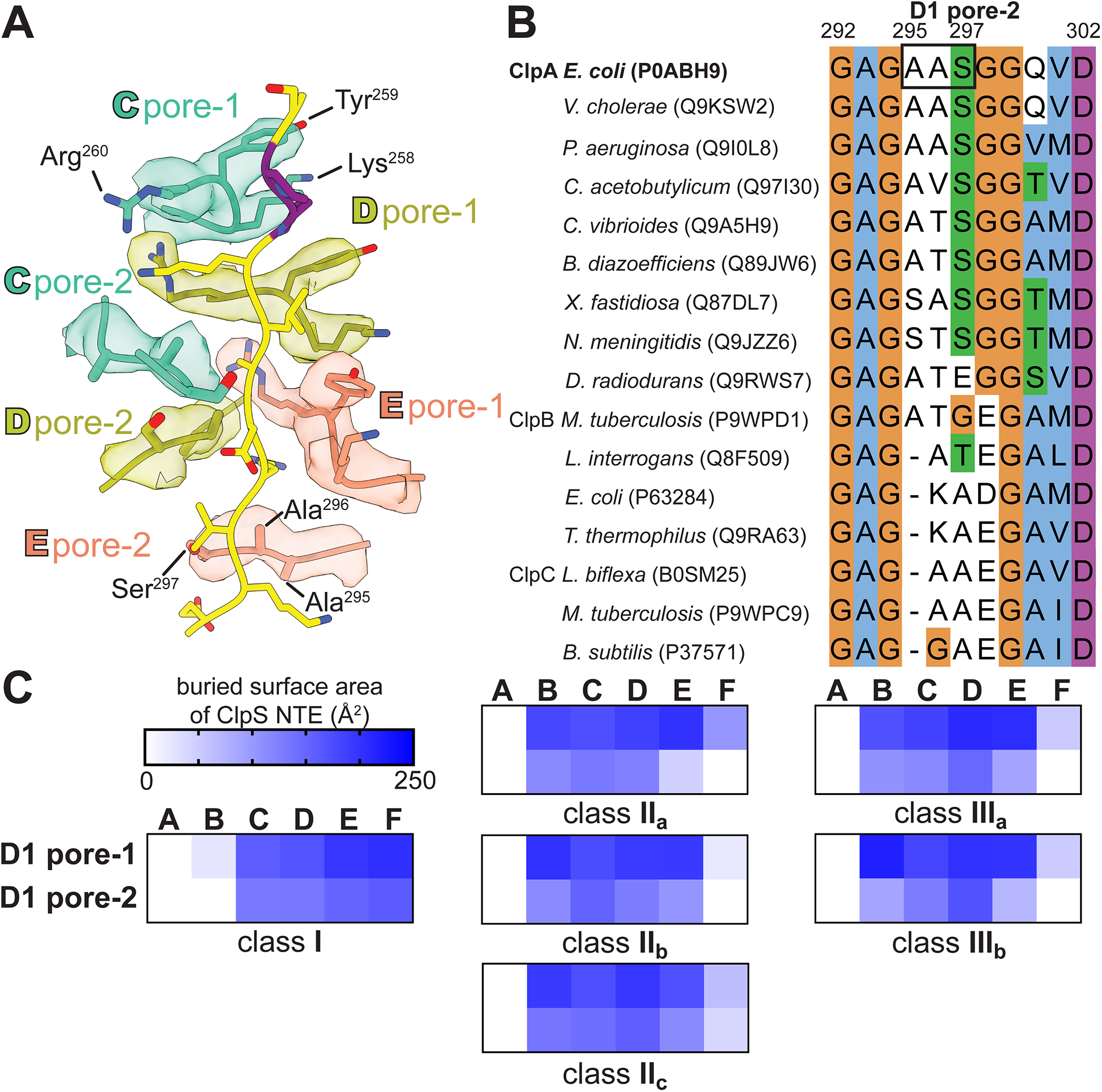
Interaction of ClpA pore-2 loops with the ClpS NTE in the D1 ring. (A) Three pairs of pore-1 (K<u>Y</u>R, res. 258–260) and pore-2 (AAS, res. 295–297) loops in the ClpA D1 ring of class **I**, shown as sticks in representative subunit coloring, and the ClpS NTE (yellow and purple sticks). Cryo-EM density of each pore loop is shown with the respective transparent surfaces. (B) Multiple sequence alignment of ClpABC family members corresponding to D1 pore-2 loops of *E. coli* ClpA (res. 292–302). UniProt accession numbers are listed in parentheses. The alignment at each position is colored according to ClustalX (orange=Gly, blue=hydrophobic, green=polar, magenta/purple=positive charge, white=unconserved). (C) Buried surface area of the ClpS NTE by the pore-1 or pore-2 loops of the D1 ring in the atomic models of classes **I**, **II**, and **III**. See also Figure S7.

To quantify the extent of pore-1 vs. pore-2 loop interactions in the D1 ring, we calculated the buried surface area (BSA) of the ClpS–NTE interface with each class of pore loops using PISA (Krissinel and Henrick, 2007). Mirroring the pattern of D1 pore-1 loops bound to the ClpS NTE, multiple pore-2 loops made significant ClpS NTE interactions in all class **I**, **II**, and **III** structures (**Figure 4C**). The D1 pore-2 loops made substantially larger contributions to the interface with the ClpS NTE than the pore-2 loops of the D2 ring, as the buried surface area contributed by the D1 pore-2 loops was comparable to those from either the D1 K<u>Y</u>R or D2 G<u>Y</u>VG pore-1 loops (**Figure S6B**). For example, in class **II**_**c**_, the BSA values for the D1 K<u>Y</u>R loops range from 63 to 196 Å^2^, the D1 pore-2 loops range from 40 to 156 Å^2^, and D2 G<u>Y</u>VG pore-1 loops range from 101 to 179 Å^2^. In contrast, the D2 pore-2 loops only weakly contacted the NTE, as BSA values of these interactions range from 20 to 74 Å^2^. Thus, pore loops in the D1 ring make a greater total number of NTE interactions than pore loops in the D2 ring. The extensive network of NTE-engaging residues in the D1 ring suggests that it has more specific polypeptide binding/recognition ‘capacity’ than the D2 ring, as predicted by biochemical studies (Hinnerwisch et al., 2005a; Zuromski et al., 2021).

### D1 pore-2 loops mediate substrate unfolding and mechanical remodeling of ClpS

To test the functional importance of the D1 pore-2 loops, we mutated the AAS sequence (res. 295–297) to increase bulkiness (QTQ), to mimic the pore-1 loop (K<u>Y</u>R), to increase flexibility (GGG), or to delete this tripeptide (Δ295–297). As a defect in ClpS binding with these mutants was one reasonable hypothesis based on our structures, we first assayed assembly of ternary ClpA_6_•ClpS•N-degron peptide complexes using fluorescence anisotropy (**Figure 5A**). Strikingly, all pore-2 loop variants maintained tight affinity for the ClpS•N-degron complexes and behaved similarly to wild-type ClpA (^WT^ClpA) in the control experiment that monitored the binary affinity of ClpA to the N-degron peptide. We then used these variants to assay ClpAPS degradation of the N-degron substrate YLFVQELA-GFP (**Figure 5B**). Notably, all of the D1 pore-2-loop variants except QTQ were unable to degrade this substrate. These defects could arise from an inability to unfold or translocate YLFVQELA-GFP or from failure to transfer the YLFVQELA-GFP substrate from ClpS to ClpA.

**Figure 5.**
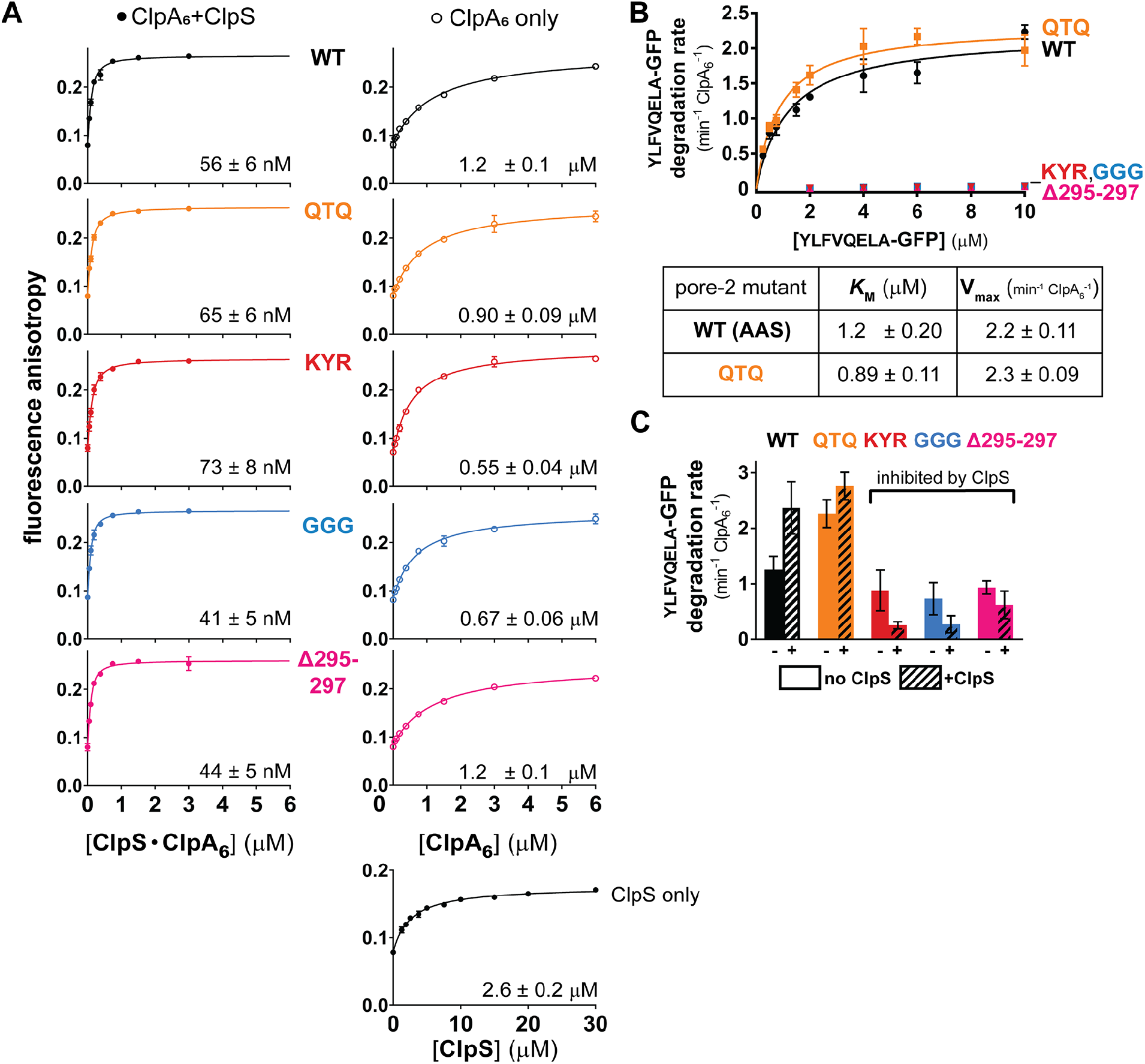
D1 pore-2 loops are critical for ClpS-mediated degradation. (A) Fluorescence anisotropy of ClpA pore-2 variants alone (ClpA_6_, open circles), with an equimolar mixture with ClpS (+ClpS, filled circles), or ClpS only (bottom right panel), titrated in increasing concentrations against a fixed concentration of fluorescein-labeled N-degron peptide (100 nM LLYVQRDSKEC-fl*) in the presence of 2 mM ATPγS. Values are mean fluorescence anisotropy values from triplicates with error bars representing ± 1 S.D, and were fit to equations listed in the *Methods*. The K_d_ values are reported on the lower right on each isotherm, with (±) the standard error of nonlinear least-squares and R^2^ values=0.99 for all fits. (B) Kinetic analysis of YLFVQELA-GFP degradation by ClpA D1 pore-2 variants (see *Methods* for concentrations). Values are mean degradation rates (min^−1^ ClpA_6_^−1^) of triplicates with error bars representing ± 1 S.D. In the table, *K*_M_ and V_max_ values ± errors were obtained by non-linear least squares fitting to the Michaelis-Menten equation. Degradation rates of GGG, Δ295–297, and K<u>Y</u>R could not be fit to the Michaelis-Menten equation. (C) YLFVQELA-GFP (20 μM) degradation rates of ClpA D1 pore-2 variants (0.1 μM ClpA_6_) and ClpP (0.2 μM ClpP_14_), in the absence and presence of ClpS (0.6 μM ClpS). Summary data are mean degradation rates (min^−1^ ClpA_6_^−1^) of triplicates with error bars representing ± 1 S.D.

For each variant, we then determined the ATP-hydrolysis rate of ClpA alone and in the presence of ClpP, ClpS, and/or a directly recognized protein substrate. The ATPase rate serves as an indirect readout of functional ClpA assembly with its binding partners, which differentially modulate ATP hydrolysis by ClpA. For instance, ClpP binding stimulates the ATPase rate of ^WT^ClpA ~two-fold, whereas ClpS suppresses the ATPase activity of ClpAP to a rate similar to that of ClpA alone (Hinnerwisch et al., 2005b; Hou et al., 2008). All of the pore-2 variants had basal ATPase rates comparable to ^WT^ClpA and exhibited ATPase modulation by ClpP and ClpS that was generally similar to wild-type (**Figure S7A**). Furthermore, in the presence of the super-folder GFP substrate (^SF^GFP-ssrA), which does not require ClpS for recognition and degradation, the ATPase rate of each ClpAP variant (with the exception of ^KYR^ClpAP) was moderately reduced during substrate processing, as expected from a previous study reporting ~20% suppression of ATP hydrolysis by GFP-ssrA (Kress et al., 2009). We conclude based on these studies that our D1 pore-2 loop mutations do not grossly alter ClpA ATPase activity and are also unlikely to substantially change ClpA assembly with ClpP, ClpS, or ^SF^GFP-ssrA.

Next, we assayed the ability of these D1 pore-2-loop variants to degrade FITC-casein, a molten-globule protein that does not require ClpS for recognition or robust ClpAP unfolding activity for degradation (Thompson et al., 1994). ^KYR^ClpAP degraded FITC-casein ~30% slower than ^WT^ClpAP, but the remaining D1 pore-2 variants degraded this substrate at roughly the wild-type rate (**Figure S7B**), indicating that recognition and translocation of this substrate are not substantially affected by the D1 pore-2 loop mutations. We then assayed the effects of the ClpA D1 pore-2 loop mutations on the steady-state kinetics of ^SF^GFP-ssrA degradation, a highly stable native substrate (**Figure S7C**). *K*_M_ values for degradation of this substrate by ^WT^ClpAP and the D1 pore-2 loop variants were within error, suggesting that the D1 pore-2 loops play little, if any, role in recognition of this substrate. V_max_ for ^SF^GFP-ssrA degradation was unaffected by the QTQ mutation, reduced ~two-fold by the GGG and Δ295–297 mutations, and reduced ~six-fold for the K<u>Y</u>R mutant. Based on these results, we conclude that the D1 pore-2 loops can promote, but are not essential for, a reaction step after initial substrate recognition, presumably GFP unfolding, which is rate limiting for degradation (Singh et al., 2000). Importantly however, these partial defects in unfolding by the GGG, Δ295–297, and KYR variants are insufficient to explain the complete inability of these mutants to degrade YLFVQELA-GFP when delivered by ClpS.

In comparison to FITC-casein and ^SF^GFP-ssrA, which are directly recognized by ClpAP, degradation of ClpS-dependent substrates require an additional protein-remodeling step. That is, ClpA must remodel ClpS, to allow substrate transfer to ClpA, and then unfold the N-degron substrate. Concurrently, ClpS reduces the ClpA ATPase rate, which in turn slows unfolding and translocation (Dougan et al., 2002; Hou et al., 2008; De Donatis et al., 2010; Román-Hernández et al., 2011; Rivera-Rivera et al., 2014; Torres-Delgado et al., 2020). Therefore, ClpS should inhibit N-degron substrate degradation by the ClpA D1 pore-2 loop variants that we infer lack sufficient unfolding activity to remodel ClpS and transfer the substrate from ClpS to ClpA. We tested this hypothesis by measuring the degradation rates of YLFVQELA-GFP in the absence and presence of ClpS (**Figure 5C**). Although recognition of N-end-rule substrates by ClpAP alone is intrinsically weak and normally enhanced by ClpS (Wang et al., 2007), the addition of ClpS hindered YLFVQELA-GFP degradation by the KYR, GGG, and Δ295–297 variants, but not by ^WT^ClpA or the QTQ variant, as predicted if the D1 pore-2 mutants are specifically defective in a ClpS remodeling step required for efficient N-degron substrate degradation.

In summary, these data suggest that the sequence identity of the AAS tripeptide (res. 295–297) alone is not critical for D1 pore-2 loop activity, as substituting these residues with QTQ had little effect on ATP hydrolysis and degradation of all substrates tested (**Figure 5B-C**; **Figure S7**). Instead, changing the chemical/conformational properties of this loop by altering charge/aromaticity (KYR) or flexibility (GGG and Δ295–297) had more profound effects. The severe defects in ClpAPS degradation conferred by the deleterious D1 pore-2 mutations but unchanged ClpS•N-degron assembly support the conclusion that D1 pore-2 loops assist in mechanical work needed to transfer the N-degron substrate from the adaptor to the protease (and perhaps also for subsequent reaction steps) but are not required for adaptor/substrate docking with ClpAP.

## DISCUSSION

### The ClpS NTE is a “degron mimic”

Our ClpAP-ClpS structures, taken with previous ClpAP structures and those of additional AAA+ family members, illustrate the variety of functional conformations AAA+ unfoldases can adopt to perform their biological functions. Importantly in all our structures, interactions between the ClpS NTE and pore loops in the ClpA channel mimic contacts observed with a polypeptide segment of the protein substrate in prior ClpA structures (Lopez et al., 2020). Specifically, the conserved tyrosines from adjacent pore-1 loops in the D1 (K<u>Y</u>R) and D2 (G<u>Y</u>VG) ring contact every second residue of the NTE polypeptide (**Figure S5**), with additional contacts mediated by the pore-2 loops of the D1 ring. Thus, in addition to its interaction with the ClpA N-domain, ClpS uses its NTE to dock tightly with the ClpA channel during substrate delivery.

Our ClpAPS structures were assembled in ATPγS, which ClpA does not hydrolyze, demonstrating that binding of the entire ClpS NTE within both ClpA rings does not require hydrolysis-dependent power strokes. Together with biochemical studies and structures of substrate complexes with ATPγS-bound ClpA (Hoskins et al., 1998, 2000; Lopez et al., 2020), these results suggest that any polypeptide in an unfolded/misfolded protein could passively enter an open ClpA channel, enabling ClpAP to function broadly in general protein quality control. Indeed, most of the ClpS NTE sequence is poorly conserved among orthologs and can be changed without compromising delivery of N-end-rule substrates (Hou et al., 2008), suggesting that ClpA can engage many different sequences. By contrast, the full axial channel of the ClpXP protease is blocked by a pore-2 loop prior to initiation of unfolding and translocation, probably limiting binding to proteins bearing highly specific ClpX degrons (Fei et al., 2020b).

Our structures also reveal that ClpA pore loops bind and engage the ClpS NTE, and thus can apply mechanical force to the ClpS core domain during the N-degron delivery process. Previous studies demonstrate that the ClpS NTE enters the ClpA axial channel during assembly of delivery complexes and also can independently function as a degron for ClpAP, providing biochemical evidence that the ClpA pore loops can ‘pull’ on the NTE to remodel ClpS (Román-Hernández et al., 2011; Rivera-Rivera et al., 2014). Blocking the ClpS NTE from entering the channel inhibits ClpS-assisted substrate degradation, reinforcing the importance of ClpA ‘pulling’ on the ClpS NTE during N-degron delivery (Rivera-Rivera et al., 2014). The degron-like binding of the NTE provides a structural basis for the delivery mechanism depicted in **Figure 6**, in which ClpA pore loops engage the NTE and power strokes resulting from ATP hydrolysis transmit force to mechanically remodel ClpS and thereby promote transfer of the N-end-rule substrate from ClpS to ClpAP for degradation.

**Figure 6.**
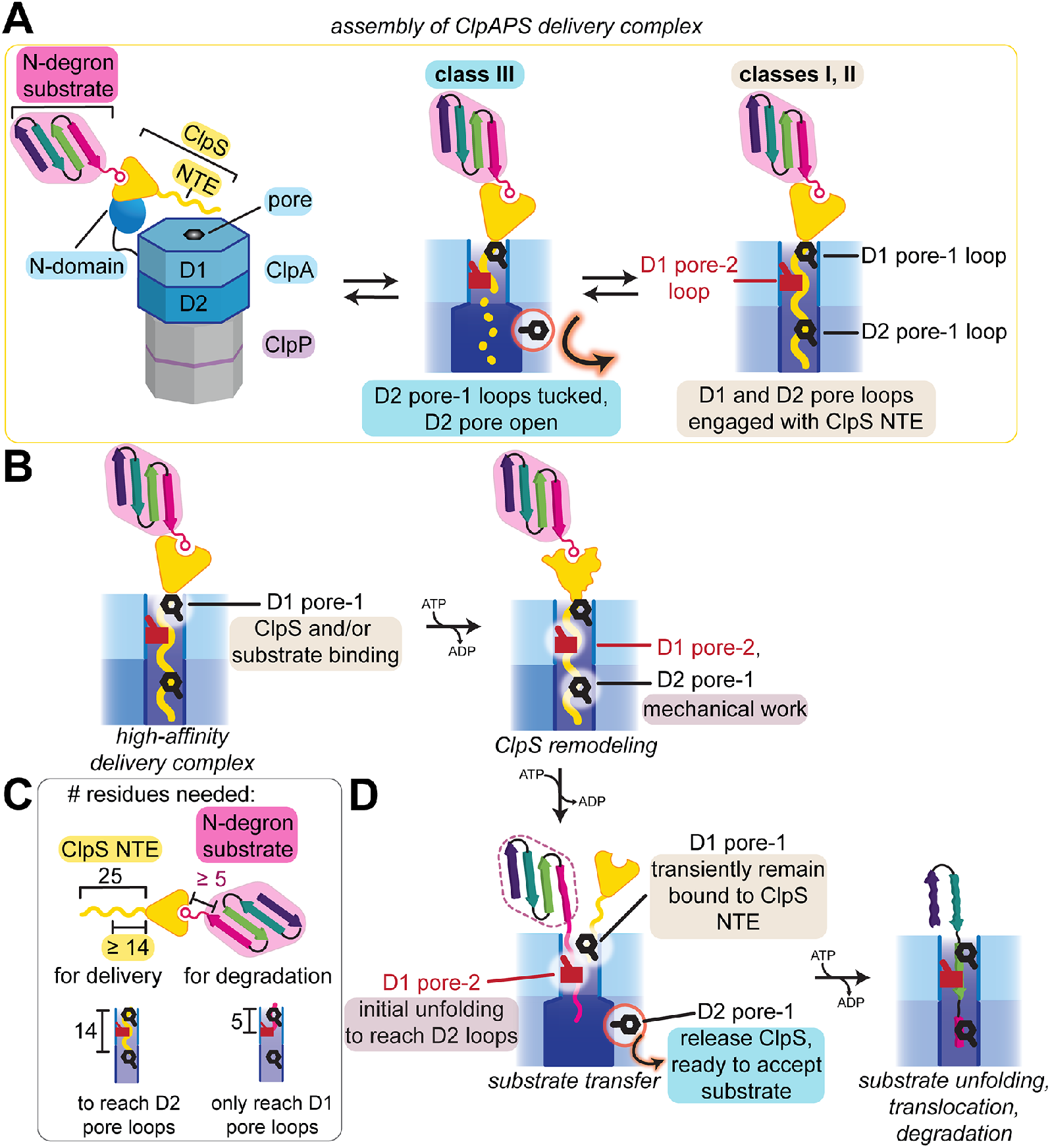
Models of ClpS-mediated degradation by ClpAP. (A) Summary of observed interactions of ClpS NTE and pore-1 and pore-2 loops in D1 and D2 from cryo-EM structures. In a dynamic equilibrium of low-affinity (not observed in this study) and high-affinity delivery complexes (classes **I, II**, and **III**), pore-1 loops in the D2 ring (*i*) are tucked-in and turned away from the axial channel (class **III**), correlated with a loss of observed ClpS NTE density, as indicated by the dotted line or (*ii*) contact the ClpS NTE (classes **I**, **II**). (B) Pore loop functions in ClpS remodeling. Pore-1 loops, especially in D1, are required for ClpS and substrate binding to form delivery complexes. ATP hydrolysis powers ClpS remodeling, allowing D1 pore-2 and D2 pore-1 loops to translocate and tug on the ClpS NTE to promote substrate transfer and degradation. (C) Summary from biochemical studies (Erbse et al., 2006; Wang et al., 2008b) of number of residues required for ClpS NTE and linker between N-end residue and folded domain in N-degron substrate. (D) Proposed function of pore loops during substrate transfer. Following ClpS remodeling, D2 pore-1 loops may release the ClpS NTE, allowing for unfolding and translocation of the N-degron substrate to proceed from the D1 pore-2 loops.

Despite the degron-like interactions of the NTE with ClpA, ClpS is not degraded (Dougan et al., 2002; Román-Hernández et al., 2011). Interestingly, mutation of Pro^24^-Pro^25^ to Ala^24^-Ala^25^ near the ClpS NTE-core junction generates a ClpS variant that can be degraded by ClpAP (Rivera-Rivera, 2015; Zuromski et al., 2021). In our structures, Pro^24^-Pro^25^ binds near the top of the ClpA channel, and the ClpS core domain is flexibly positioned directly above the pore. Although our structures only capture the initial docking of ClpS with ClpA, we propose that during subsequent stages of ClpS delivery, ClpA pore loops may not grip Pro^24^-Pro^25^ strongly enough to fully unfold ClpS, leading to ‘back-slipping’ in the channel and thus ClpS release. Such slipping is likely a consequence of the unique chemical properties of proline, which lacks an amide hydrogen and cannot form the extended peptide conformation adopted by the rest of the NTE in our structures and by substrate polypeptides in the channels of many other AAA+ unfoldases (Puchades et al., 2020). ClpXP translocates poly-proline at an even slower rate and with a higher cost of ATP hydrolysis cycles than poly-glycine, another homopolymer that leads to pore-loop ‘slipping’ (Barkow et al., 2009; Bell et al., 2019). Other types of ‘slippery’ sequences adjacent to folded domains have been shown to cause release of truncated degradation products by a number of AAA+ proteases (Lin and Ghosh, 1996; Levitskaya et al., 1997; Sharipo et al., 2001; Hoyt et al., 2006; Daskalogianni et al., 2008; Kraut et al., 2012; Kraut, 2013; Too et al., 2013; Vass and Chien, 2013; Bell et al., 2019). Partial ClpAP processing of native ClpS does not occur because its NTE is not long enough to enter the ClpP peptidase chamber (**Figure 1F**), but is observed for a variant bearing a duplicated NTE of ~50 residues (Rivera-Rivera et al., 2014).

### Implications of tucked pore-1 loops in the D2 ring

In the D2 ring of our class**-III** structures, many G<u>Y</u>VG pore-1 loops assume a ‘tucked’ conformation in which they rotate away from the center of the ClpA axial channel and do not engage the ClpS NTE (**Figure 3B-C**), presumably weakening ClpS•ClpA binding. There are several functional implications. First, the D2-disengaged/D1-engaged species could represent an intermediate in the assembly of higher-affinity ClpAPS complexes in which both rings engage the NTE (**Figure 6A**). Second, tucked D2 G<u>Y</u>VG pore loops could be important during latter steps in ClpS-dependent substrate delivery (**Figure 6B**), which require conformational remodeling of the ClpS core to weaken its interactions with the N-degron substrate and promote its transfer to ClpA (Hou et al., 2008; Román-Hernández et al., 2011; Rivera-Rivera et al., 2014). For example, after failed attempts by ClpA to fully unfold the ClpS core, release of the NTE from the D2 ring could increase the probability that ClpS dissociates completely from ClpAP, freeing the D2 pore loops to engage the N-degron substrate for degradation (see **Figure 6D**). Finally, pore-loop tucking does not require ATP hydrolysis, suggesting that under certain conditions (*e.g*., when bound to the ClpS NTE), ClpA readily adopts the class-**III** structures, which constitute ~20% of particles in our final dataset.

More broadly, pore-loop tucking may be used during the process of enzyme pausing and/or unloading by ClpA and other AAA+ unfoldase motors. For example, the ClpA D1 ring functions as a ‘back up’ motor to prevent pausing when the principal D2-ring motor fails (Kotamarthi et al., 2020; Zuromski et al., 2021). Transiently breaking contacts with the polypeptide via pore-loop tucking in only the D2 ring would allow the weaker D1 ring to continue unfolding/translocation without working against the stalled D2 motor. Subsequent ‘untucking’ of the D2 pore-1 loops once the sequence causing the pause is cleared would allow the D2 motor to re-engage, restarting robust translocation by both rings. More generally, concerted loss of peptide contacts by all AAA+ domains within a ring via pore-loop rotation and tucking would be an efficient mechanism for an unfoldase either to transiently disengage from a bound polypeptide or facilitate full enzyme dissociation upon failure of a AAA+ motor to unfold, translocate, or remodel a bound protein. By contrast, dissociation of a AAA+ enzyme from its polypeptide track by transitioning from a closed, substrate-bound right-handed spiral to an open, left-handed ‘lock-washer’ observed in some Hsp100 family members (Yokom et al., 2016; Gates et al., 2017; Yu et al., 2018) requires much larger, global conformational changes throughout the AAA+ hexamer.

### Specialized functions of pore-2 loops

The pore-2 loops of other AAA+ unfoldases/remodeling enzymes have been shown to contact substrate polypeptides (Johjima et al., 2015; Deville et al., 2017; Gates et al., 2017; Puchades et al., 2017, 2019; Alfieri et al., 2018; Yu et al., 2018; Sandate et al., 2019; Zehr et al., 2020; Fei et al., 2020a, 2020b; Han et al., 2020; Shin et al., 2021; Kavalchuk et al., 2022). The AAS residues of the ClpA D1 pore-2 loops make substantial contacts with the ClpS NTE. Nevertheless, we find that these interactions are not critical for ClpA•ClpS binding but instead help mediate mechanical work needed during ClpS-assisted N-end-rule degradation (**Figure 5; Figure 6B**). We propose that the D1 pore-2 loops of ClpA collaborate with the D2 pore-1 loops, which are also required for ClpS delivery (Zuromski et al., 2021), in mechanical remodeling of ClpS and/or substrate transfer to ClpA. Both sets of loops could contribute to coordinated pulling on the NTE to apply force to and remodel ClpS. Next, the pore-2 loops could capture and initiate unfolding of the ‘released’ N-degron substrate, and generate a sufficiently long polypeptide ‘tail’ to reach the more powerful D2 pore-1 loops (**Figure 6C-D**). Meanwhile, the D2 pore-1 loops could release the ClpS NTE via concerted loop-tucking, but then ‘untuck’ to grab this ‘tail’ for processive substrate unfolding and translocation. Future studies parsing the interaction of pore-1 and pore-2 loops in both ClpA rings are needed to further elucidate the mechanistic steps of N-degron substrate delivery and degradation, as well as to understand why pore-2 loops are critical in ClpS-mediated degradation but less important for other classes of substrates.

### ClpA functions using both coordinated and independent action of the D1 and D2 rings

Loss of D2 pore-1 loops engagement with the ClpS NTE is a major feature distinguishing our class-**III** structures from classes **I** and **II**. Although the asymmetric engagement of substrate in the D1 but not the D2 ring of class **III** has some parallels with substrate-bound structures of NSF and Pex1•Pex6 (Blok et al., 2015; Gardner et al., 2018; White et al., 2018), the substrate-binding rings of these other enzymes adopt a ‘canonical’ right-hand spiral organization, whereas the ring that does not bind substrate assumes a planar conformation. In contrast, the non-binding, ClpA D2 ring in class **III** remains in the right-handed spiral conformation. Moreover, the portion of the ClpS NTE in the D1 ring of class-**III** structures is bound in the same fashion as our class-**I** and class-**II** structures. This structural snapshot of ‘divided’ NTE engagement between the D1 and D2 pore-1 loops reinforces biophysical and biochemical experiments that reveal a division of labor between the two AAA+ modules of ClpA (Kress et al., 2009; Kotamarthi et al., 2020; Zuromski et al., 2021). Multiple studies of other double-ring remodeling/unfoldase enzymes, including ClpB, Hsp104, ClpC, Cdc48/p97/VCP, and the ribosomal assembly factor Rix7, report the separation of substrate binding/recognition functions in one ring from the role of the second ring as the principal motor performing mechanical work (Hattendorf and Lindquist, 2002; Mogk et al., 2003; Wang et al., 2011; Doyle et al., 2012; Bodnar and Rapoport, 2017a, 2017b; Lo et al., 2019). These results illustrate that functional specialization of individual rings is emerging as a theme shared by many double-ring AAA+ unfoldases and protein-remodeling enzymes.

## Author Contributions

Conceptualization, S.K., X.F., R.T.S., and T.A.B.; Methodology, S.K., X.F., T.A.B., and R.T.S; Validation, S.K., X.F., and R.T.S.; Formal Analysis, S.K. and X.F.; Investigation, S.K. and X.F.; Resources, S.K.; Writing – Original Draft, S.K. and X.F.; Writing – Review & Editing, S.K., X.F., R.T.S., and T.A.B.; Visualization, S.K. and X.F.; Supervision, R.T.S. and T.A.B.; Funding Acquisition, S.K., R.T.S., and T.A.B.

## Acknowledgments

We thank E. Brignole and P. Dip for support in preparing and screening cryo-EM grids at the MIT.nano Automated Cryogenic Electric Microscopy Facility on a Talos Arctica microscope, which was a gift from the Arnold and Mabel Beckman Foundation, and C. Xu, K. Song, and K. Lee for data collection at the Cryo-EM Core Facility at the University of Massachusetts Chan Medical School. We thank I. Levchenko for advice preparing ClpAPS complexes and S. Bell, T. Bell, J. Park Morehouse, T. Shih, J. Zhang, and K. Zuromski for helpful advice and feedback.

This work was supported by NIH grant AI-016892 (R.T.S., T.A.B.), the Howard Hughes Medical Institute (T.A.B.), and the National Science Foundation Graduate Research Fellowship grant 1745302 (S.K.). The content is solely the responsibility of the authors and does not necessarily represent the official views of the National Institutes of Health.

## Declarations of Interest

The authors declare no competing interests.

---

**Figure S1.**
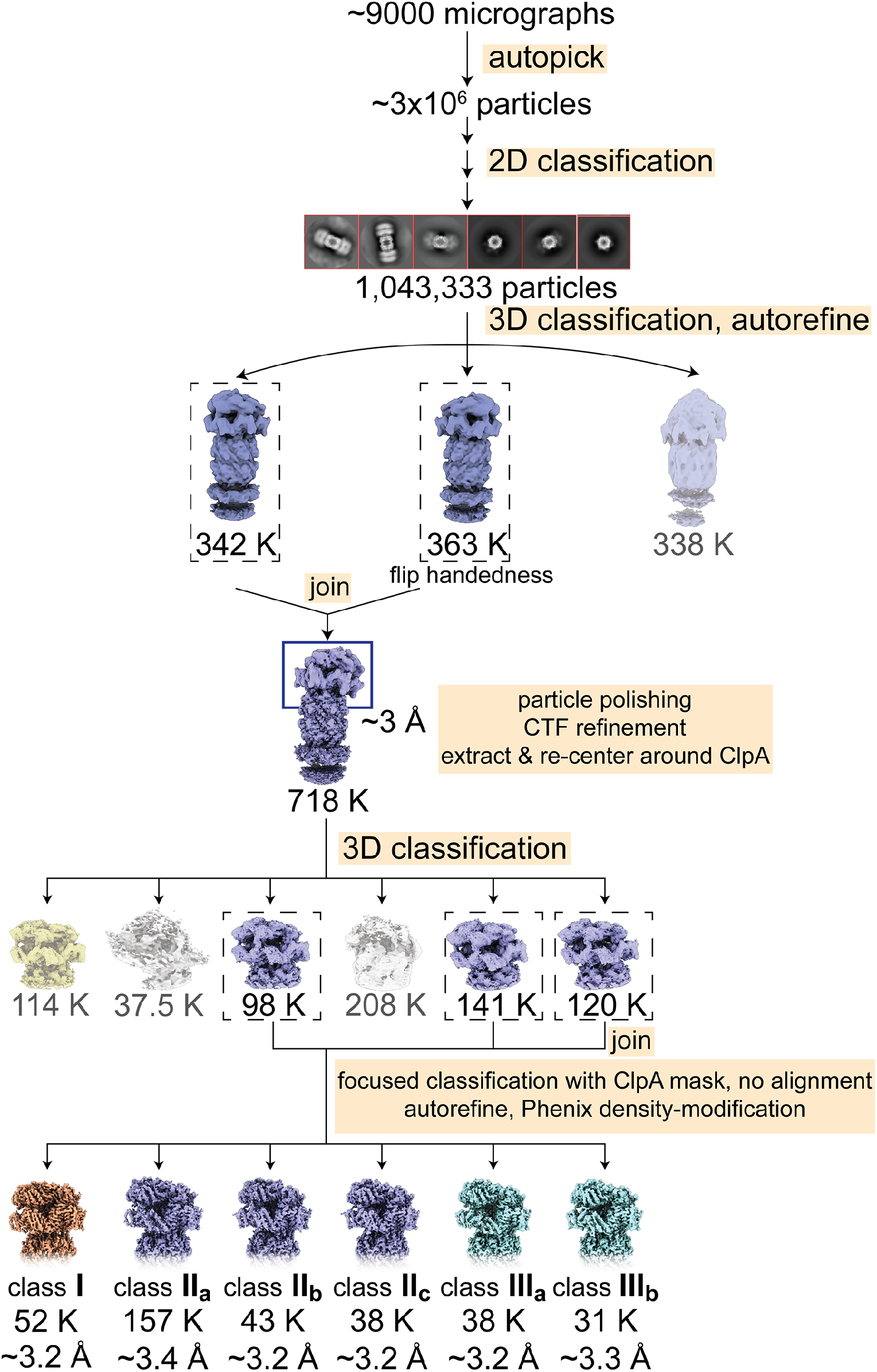
Cryo-EM data-processing workflow diagram, Related to Figure 1. EM micrographs containing doubly-capped ClpAP complexes (two ClpA hexamers bound to one ClpP 14-mer) were processed in RELION-3. The final 3D classes were refined using density-modification in PHENIX.

**Figure S2.**
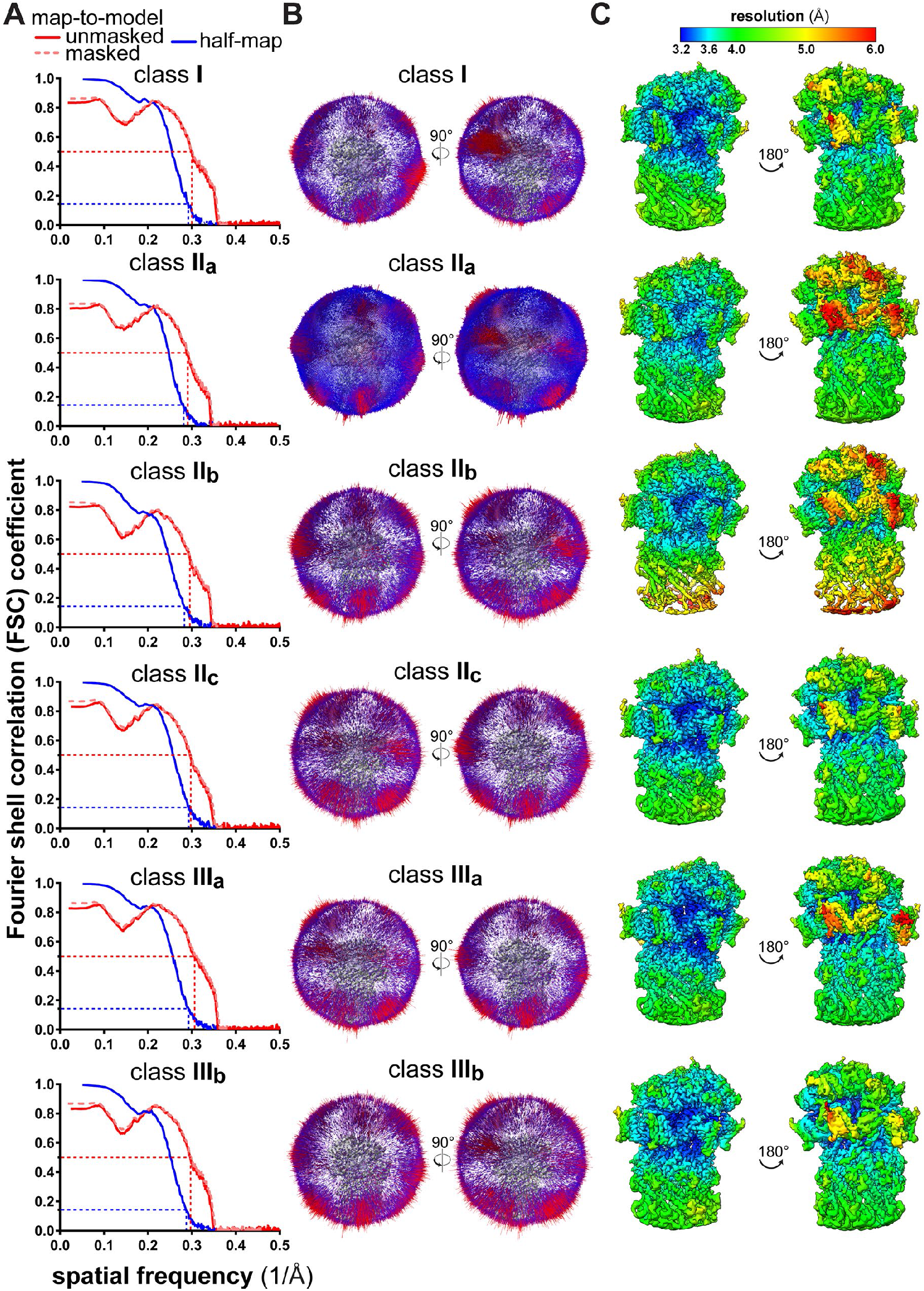
Cryo-EM validation, Related to Figure 1. (A) Fourier Shell Correlation (FSC) plots of half maps (shown in blue) or model-map (shown in red) resolution. The dashed lines indicate the cut-off values at FSC=0.5 (model-map) or FSC=0.143 (half-map). (B) Euler angle distribution plots of the particles used in the final reconstruction of class **I**, **II**_**a-c**_, and **III**_**a,b**_ structures. (C) Local resolution maps of final reconstructions, colored according to RELION-3 calculations.

**Figure S3.**
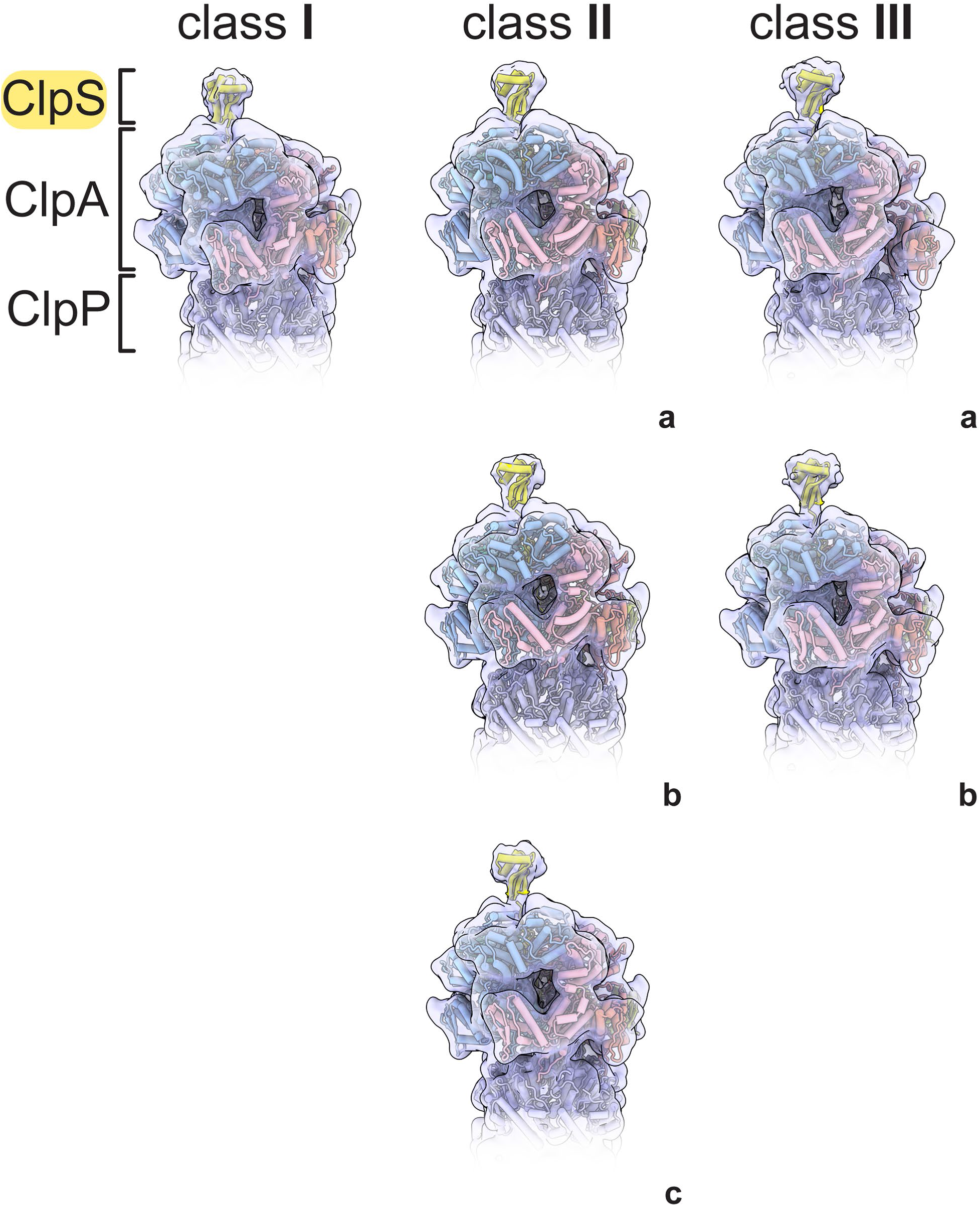
ClpS docking to cryo-EM maps, Related to Figure 1. The ClpS core domain (PDB 3O2B; res. 27-106) was docked to final reconstructions of class **I**, **II**_**a-c**_, and **III**_**a,b**_ that were low-pass filtered to 10 Å.

**Figure S4.**
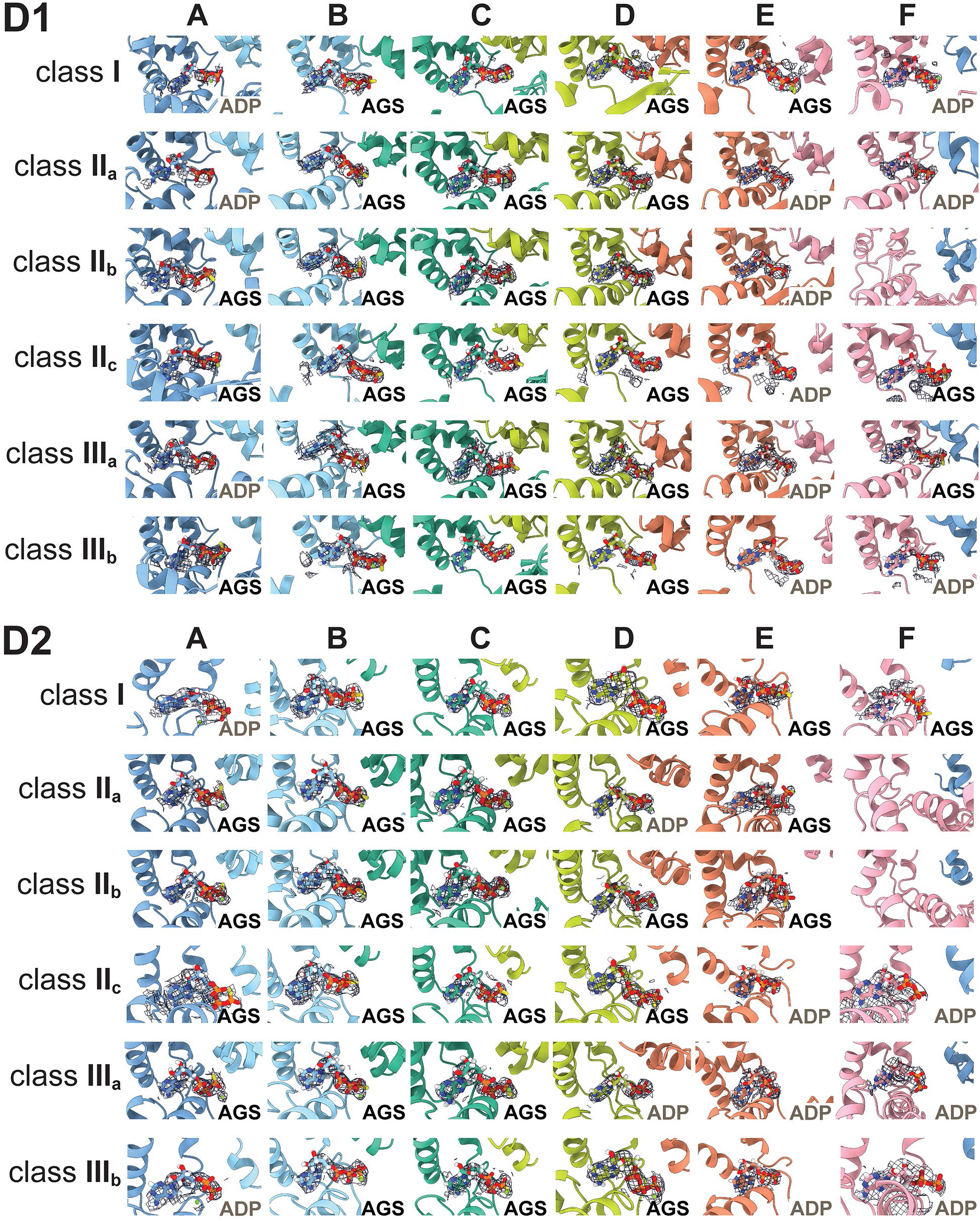
Nucleotide occupancy, Related to Figure 2, Related to Figure 3. EM density (shown as mesh) of nucleotides in each D1 and D2 binding site of class **I**, **II**_**a-c**_, and **III**_**a,b**_ structures. Nucleotide density is not observed in the D1 site of F subunit in **II**_**b**_ or the D2 site of the F subunit of **II**_**a**_ or **II**_**b**_.

**Figure S5.**
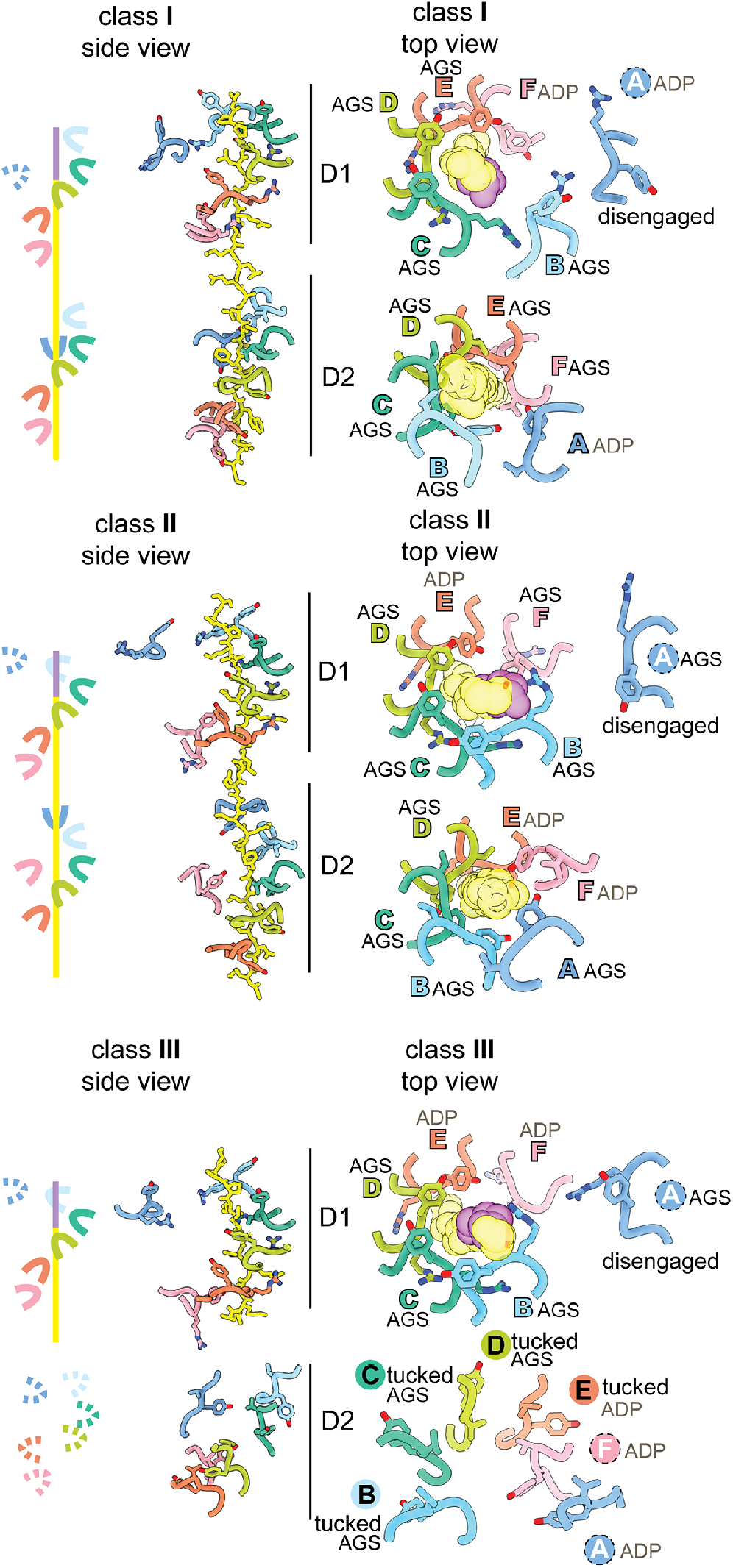
Pore-1 loop interactions with ClpS NTE, Related to Figure 3B-C. The leftmost panel in each structure is a diagram of pore-1 loops in the D1 and D2 rings, where solid lines represent the presence of NTE contacts and dashed lines represent their absence. Middle panel shows lateral views of contacts between ClpA pore-1 loops and the ClpS NTE in class **I**, **II**_**c**_, and **III**_**b**_ atomic models. The rightmost panel in each structure is a closer view of the pore-1 loops with the NTE shown as transparent spheres and the corresponding nucleotide from each subunit. Labels in colored text denote NTE engagement; the dotted circle denotes lack of NTE engagement, with Tyr^540^ pointing towards the channel; labels in black text indicate the tucked conformation (Tyr^540^ away from the channel).

**Figure S6.**
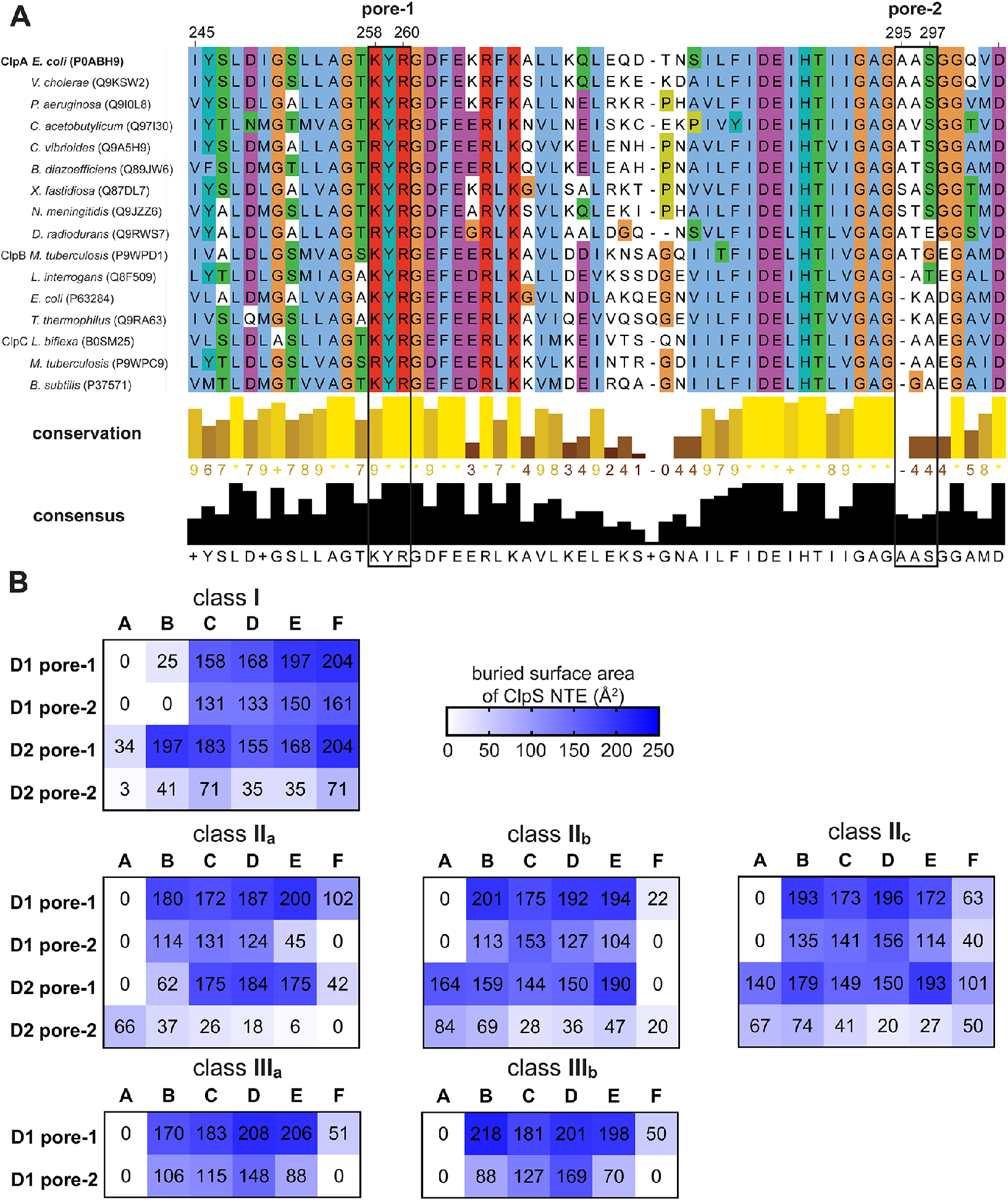
Comparison of pore-1 and pore-2 loop sequences and contacts to ClpS NTE, Related to Figure 4B-C. (A) Multiple sequence alignment of ClpABC family members, corresponding to pore-1 (res. 258–260, *E. coli* ClpA) and D1 pore-2 loops of *E. coli* ClpA (res. 292–302), created using MUSCLE alignment (Edgar, 2004). UniProt accession numbers are listed in parentheses. The alignment at each position is colored according to ClustalX (orange=Gly, blue=hydrophobic, green=polar, magenta/purple=positive charge, white=unconserved). Conservation scores are calculated in Jalview from the amino acid properties in the alignment. Conserved columns that have the highest conservation score are indicated by ‘*’ symbols (corresponding to a numeric score of 11), followed next by mutations that conserve all physico-chemical properties, indicated by ‘+’ symbols. Gaps are indicated by ‘-’, and the lowest conservation score is zero. (B) Buried Surface Area (BSA) of ClpS NTE. Contacts between ClpS NTE and pore-1 or pore-2 loops in the D1 and D2 rings of classes **I**, **II**_**a-c**_, and **III**_**a,b**_ were evaluated using PISA. Raw BSA values are provided in each box that correspond to coloring by the heat map scale.

**Figure S7.**
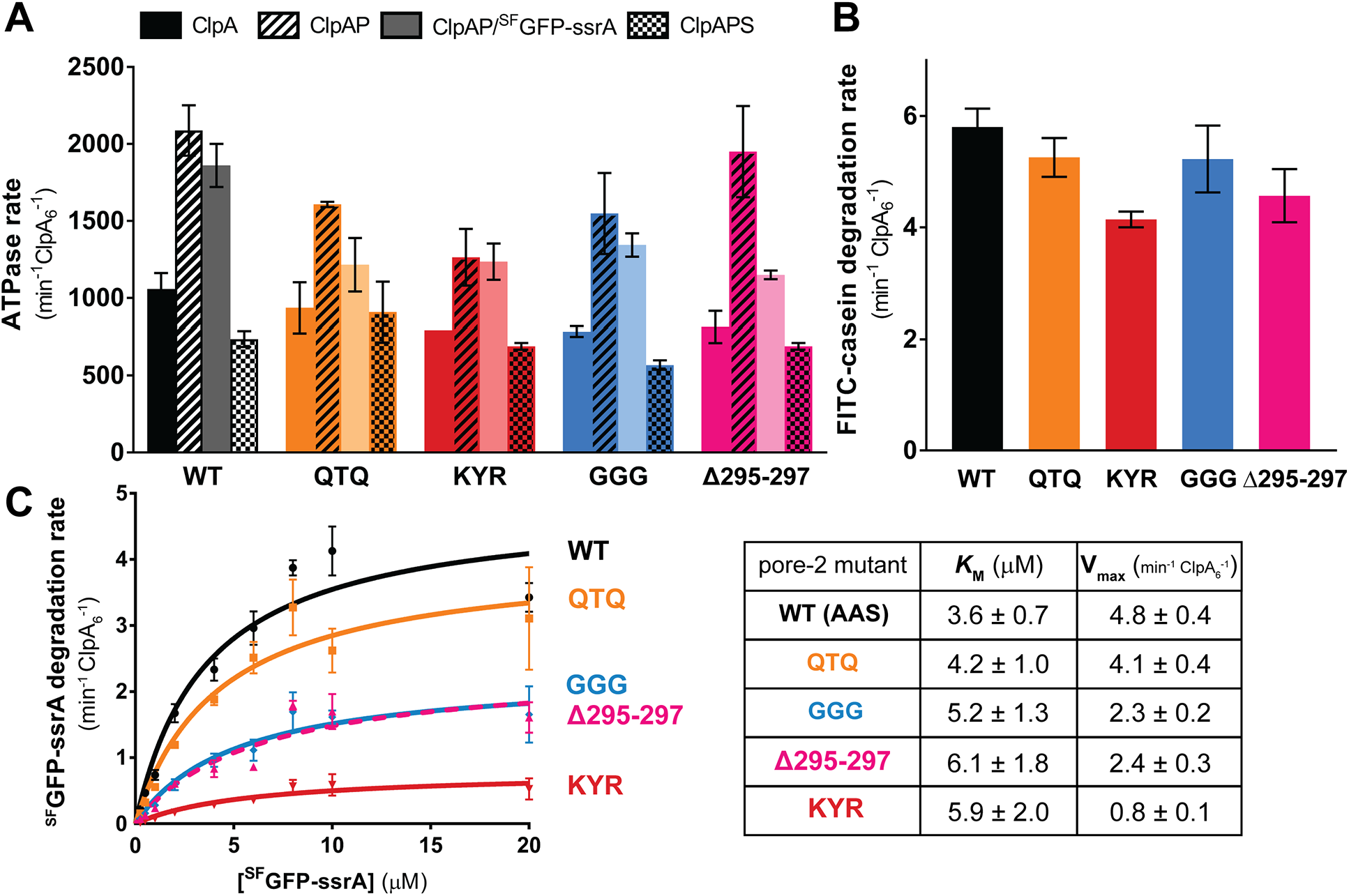
D1 pore-2 loop variants can support ATP hydrolysis and substrate unfolding, translocation, and degradation, Related to Figure 5. (A) ATP hydrolysis rates of ClpA D1 pore-2 variants alone or in the presence of ClpP, ClpP and ^SF^GFP-ssrA substrate, or ClpP and ClpS; see *Methods* for concentrations. (B) FITC-casein (20 μM) degradation rates of ClpA D1 pore-2 variants (0.2 μM ClpA_6_) and ClpP (0.4 μM ClpP_14_). Summary data are mean degradation rates (min^−1^ ClpA_6_^−1^) of triplicates with error bars representing ± 1 S.D. (C) Kinetic analysis of ^SF^GFP-ssrA degradation by ClpA D1 pore-2 variants (0.2 μM ClpA_6_, 0.4 μM ClpP_14_). Values are mean degradation rates (min^−1^ ClpA_6_^−1^) of triplicates with error bars representing ± 1 S.D. In the table, *K*_M_ and V_max_ values ± errors were obtained by non-linear least squares fitting to the Michaelis-Menten equation.

## METHODS

### Proteins and peptides

ClpA pore-2 mutations were introduced using round-the-horn mutagenesis with T4 polynucleotide kinase and Q5 high-fidelity polymerase (New England Biolabs) into pET9a-^M169T^ClpA, the plasmid used to express ClpA (gift from J. Flanagan, Hershey Medical Center). The M169T substitution helps overexpression of full-length ClpA (Seol et al., 1995) and is present in our lab version of ‘wild-type’ ClpA. ClpA and pore-2 variants were purified as described (Hou et al., 2008) and stored in HO buffer (50 mM HEPES-KOH [pH 7.5], 300 mM NaCl, 20 mM MgCl_2_, 10% [w/v] glycerol, 0.5 mM DTT). ClpP-His_6_ was expressed in *E. coli* strain JK10 (*clpP∷cat*, Δ*lon*, *slyD∷kan*, λDE3 (Kenniston et al., 2003)), purified as described (Kim et al., 2000), and stored in 50 mM Tris-HCl (pH 8), 150 mM KCl, 10% glycerol, 0.5 mM EDTA, and 1 mM DTT. ClpS and YLFVQELA-GFP were expressed in *E. coli* strain BL21(DE3) and purified as described for His_6_-SUMO-fusion proteins (Hou et al., 2008; Román-Hernández et al., 2011). ^SF^GFP-ssrA was expressed and purified as described (Nager et al., 2011). ClpS, ^SF^GFP-ssrA, and YLFVQELA-GFP were stored in 25 mM HEPES-KOH (pH 7.5), 150 mM KCl, 10% glycerol, and 1 mM DTT. FITC-casein (Sigma-Aldrich C0528) was dissolved in HO buffer and used freshly for biochemical assays; an extinction coefficient at 280 nm (11,460 M^−1^cm^−1^) and absorbance values at 280 nm and 494 nm (to calculate and correct for overlap from the fluorescence of the FITC moiety) were used to calculate its concentration. The LLYVQRDSKEC-fluorescein N-degron synthetic peptide (21st Century Biochemicals [Marlborough, MA], molecular weight 1779.9 g/mol) was dissolved and stored at 100 μM in 15% DMSO.

### Sample preparation and EM data acquisition

ClpA_6_ (4 μM), ClpP_14_ (8 μM), ClpS (13 μM), and YLFVQELA-GFP (13 μM) were mixed in 70 μL of assembly buffer (50 mM HEPES-KOH, pH 7.5, 300 mM KCl, 5 mM MgCl_2_, 2 mM TCEP, 4% glycerol, 2 mM ATPγS [Calbiochem]) for 5 min at 25 °C. 25 μL of this mixture was then chromatographed at room temperature and a flow rate of 0.04 mL/min on a Superdex-200 3.2/300 size exclusion column equilibrated in assembly buffer (GE Healthcare Ettan). A 50 μL fraction containing the largest molecular weight complex was assessed by SDS-PAGE (stained with SYPRO Red [Thermo Fisher]) and pooled for cryo-EM. After diluting the sample two-fold in assembly buffer, a 3 μL aliquot of the mixture was applied to glow-discharged R1.2/1.3 300 mesh holey carbon gold grids (Quantifoil). After a 15 s incubation, grids were blotted for 4 s at 4 °C, 100 % humidity, using Whatman grade 595 filter paper, and plunged into liquid ethane using a Vitrobot Mark IV system (Thermo Fisher Scientific).

A single grid was imaged for data collection using a Talos Arctica with a Gatan K3 direct electron detector (University of Massachusetts Chan Medical School Cryo-EM Microscopy Facility, Worcester, MA) in super-resolution mode, operated at 200 keV to collect high-resolution movies at (0.435 Å per pixel; uncalibrated magnification 45,000X) with a defocus range of −0.5 μm to −2.5 μm, with a total dose of 34.71 e^−^/Å^2^ over 26 frames (200 ms per frame).

### Cryo-EM data processing

Each movie was binned by a factor of 2, aligned, corrected for beam-induced motion using MotionCor2 (Zheng et al., 2017), and CTF estimation was calculated by CTFFIND4 (Rohou and Grigorieff, 2015). A total of 9,169 micrographs were analyzed using RELION 3.0.8 (Zivanov et al., 2018) for data processing, classification, and 3D reconstruction. The majority of auto-picked particles were doubly capped complexes consisting of two ClpA hexamers per ClpP 14-mer. Following three rounds of 2D classification, 1,043,033 particles were used for 3D reconstruction. The cryo-EM map of ClpXP (EMD-20406), which was collected under similar parameters as our dataset, was low-pass filtered to 60 Å to generate an initial model for reconstruction. After the first round of 3D classification, two of three high-quality classes were combined, totaling 717,833 particles; the second class closely resembled the first with the exception of handedness and was flipped to correct handedness before being combined. The two classes were also utilized for per-particle CTF refinement and motion correction. The combined class had a resolution of ~3 Å. The fulcrum was shifted to the center of ClpA, and particles were re-boxed to improve the resolution of this region of ClpA before performing the second round of 3D classification (T=4) with alignment to generate six classes. Three good classes were selected and combined (358,726 particles) for a third round of 3D classification with a ClpA mask, without alignment (T=20) to yield the final six classes. Each class was then subjected to 3D auto-refinement without symmetry to yield six maps with ~3.5 Å resolution. To generate the final maps, each map was density-modified and autosharpened in PHENIX (Adams et al., 2010), giving final resolutions of ranging from ~3.2–3.4 Å (**Table 1**).

### Molecular modeling and refinement

The ClpAP cryo-EM structure (PDB code 6W23) was docked into the EM map for the class-**I** structure, and the ClpAP cryo-EM structure (PDB code 6W22) was docked into all other EM maps using “fit in map” in Chimera (Pettersen et al., 2004). Real-space refinement was performed using PHENIX, and model building was performed in Coot (Emsley et al., 2004). The ClpS NTE sequence (residues 2–26) was added manually in Coot. Geometry of the final models was evaluated using MolProbity (Williams et al., 2018). Figures and movies were generated using Chimera, ChimeraX (Goddard et al., 2007), and PyMOL (Schrödinger, LLC).

### Multiple sequence alignment

The amino-acid sequences of bacterial ClpA, ClpB, and ClpC proteins were downloaded from UniProtKB (Bateman et al., 2021) and aligned using MUSCLE (Edgar, 2004) with MEGA7 (Kumar et al., 2016). The sequence alignment was visualized in Jalview (Waterhouse et al., 2009) and colored according to the Clustal X scheme.

### Buried surface area calculations

The buried surface area of the ClpS NTE in all class structures was analyzed using the ‘Protein interfaces, surfaces and assemblies’ service PISA at the European Bioinformatics Institute. (http://www.ebi.ac.uk/pdbe/prot_int/pistart.html). The BSA values were summed from the D1 pore-1 loop region (residues 254-264), D1 pore-2 loop region (residues 292-302), D2 pore-1 loop region (residues 536-544), and D2 pore-2 loop region (residues 525-531).

### Biochemical assays

Biochemical experiments were performed with at least three technical replicates at 30 °C in HO buffer using a SpectraMax M5 Microplate Reader (Molecular Devices) to measure initial rates of absorbance or fluorescence changes or equilibrium anisotropy values. ATP-hydrolysis rates were measured over the first ~2 min by monitoring the loss of absorbance at 340 nm using a coupled NADH–ATP assay (Burton et al., 2001) with 5 mM ATP (Sigma-Aldrich), pyruvate kinase (Sigma-Aldrich; P9136 at 20 units/mL), lactate dehydrogenase (Sigma-Aldrich; L1254 at 20 units/mL), 7.5 mM phosphoenolpyruvate (Sigma-Aldrich; P0564), and 0.2 mM NADH (Roche 10107735001). ATP-hydrolysis assays were performed under four conditions: (*i*) ClpA_6_ or variants (0.2 μM); (*ii*) ClpA_6_ or variants (0.1 μM) and ClpP_14_ (0.1 μM); (*iii*) ClpA_6_ or variants (0.2 μM), ClpP_14_ (0.2 μM), and ^SF^GFP-ssrA (3 μM); and (*iv*) ClpA_6_ or variants (0.2 μM), ClpP_14_ (0.2 μM), and ClpS (0.6 μM).

FITC-casein degradation assays were monitored by increases in fluorescence (excitation 340 nm, emission 520 nm) as a consequence of protease-dependent unquenching over the first 5 min; reactions contained ClpA_6_ or variants (0.2 μM), ClpP_14_ (0.4 μM), and an ATP-regeneration system (4 mM ATP, 50 μg/mL creatine kinase [Sigma-Aldrich], 5 mM creatine phosphate [Sigma-Aldrich]). The endpoint fluorescence for complete FITC-casein degradation was determined by addition of porcine elastase (100 μg/mL; Sigma-Aldrich) to each well, followed by a 30-min incubation prior to reading. To determine FITC-casein degradation rates, the increase in relative fluorescence units was normalized to the endpoint fluorescence value from fully unquenched substrate after porcine elastase incubation and the background rate was subtracted from each reaction on the basis of a no-enzyme buffer-only control.

Degradation of GFP variants was monitored by loss of fluorescence (excitation 467 nm, emission 511 nm) over the first 5-10 min. Briefly, rates were calculated by normalizing the slope values of relative fluorescence units (RFUs)/time by the fluorescence signal determined from a standard curve of RFUs versus varying concentrations of substrate in the linear range. Degradation of different concentrations of ^SF^GFP-ssrA (0.25 - 20 μM) was assayed using ClpA_6_ or variants (0.2 μM), ClpP_14_ (0.4 μM), and the ATP-regeneration system described above. Degradation of different concentrations of YLFVQELA-GFP (0.25 - 20 μM) was assayed using ClpA_6_ or variants (0.1 μM), ClpP_14_ (0.2 μM), ClpS (0.6 μM), and the ATP-regeneration system. For degradation of YLFVQELA-GFP by the GGG, K<u>Y</u>R, and Δ295-297 pore-2-loop mutants shown in Figure 5B, concentrations were ClpA_6_ variant (0.6 μM), ClpP_14_ (1.2 μM), and ClpS (3.6 μM).

The binding of the peptide LLYVQRDSKEC-fluorescein (100 nM) to (*i*) ClpA_6_ or variants (0.047 - 6 μM), (*ii)* ClpS (1.25 - 30 μM), or (*iii*) equimolar mixtures of ClpA_6_ or variants and ClpS (0.047 - 3 μM) at equilibrium was assayed by fluorescence anisotropy (excitation 490 nm, emission 525 nm) in the presence of ATPγS (2 mM). Only ClpA_6_•ClpS•peptide ternary and ClpA_6_•peptide binary complexes have higher anisotropy levels (in comparison to ClpS•peptide binary complexes) as a result of the much larger molecular weight of ClpA_6_ (~500 kDa) compared to that of ClpS (~10 kDa). Data were fit by a non-linear least-squares algorithm to equations for ClpA_6_ only and ClpS only experiments:

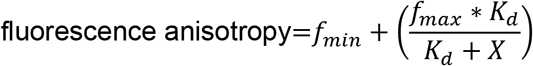

or to a quadratic equation for tight binding for ClpA_6_•ClpS complexes:

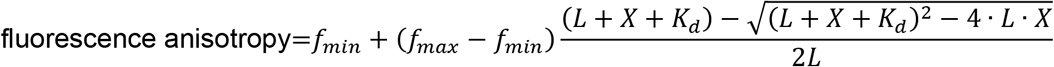

where *f*_*min*_ is the background anisotropy value, *f*_*max*_ is the maximum anisotropy value at saturated binding, *L* is the concentration of peptide (100 nM), *K*_d_ is the dissociation equilibrium constant (in nM), and *X* is the concentration of ClpA_6_•ClpS (in nM).

## Notes

### Competing Interest Statement

The authors have declared no competing interest.

